# Metabolic rules of microbial community assembly

**DOI:** 10.1101/2020.03.09.984278

**Authors:** Sylvie Estrela, Jean C. C. Vila, Nanxi Lu, Djordje Bajic, Maria Rebolleda-Gomez, Chang-Yu Chang, Alvaro Sanchez

## Abstract

To develop a quantitative theory that can predict how microbiomes assemble, and how they respond to perturbations, we must identify which descriptive features of microbial communities are reproducible and predictable, which are unpredictable, and why. The emergent metagenomic structure of communities is often quantitatively convergent in similar habitats, with highly similar fractions of the metagenome being devoted to the same metabolic pathways. By contrast, the species-level taxonomic composition is often highly variable even in replicate environments. The mechanisms behind these patterns are not yet understood. By studying the self-assembly of hundreds of communities in replicate, synthetic habitats, we show that the reproducibility of microbial community assembly reflects an emergent metabolic structure, which is quantitatively predictable from first-principles, genome-scale metabolic models. Taxonomic variability within functional groups arises through multistability in population dynamics, and the species-level community composition is predictably governed by the mutual competitive exclusion of two sub-dominant strains. Our findings provide a mechanistic bridge between microbial community structure at different levels of organization, and show that the evolutionary conservation of metabolic traits, both in terms of growth responses and niches constructed, can be leveraged to quantitatively predict the taxonomic and metabolic structure of microbial communities.

## Introduction

Understanding how the confluence of metabolic and ecological factors conspire to shape the composition and function of microbial communities is needed in order to quantitatively predict how communities assemble, and how they respond to perturbations such as diet shifts (David et al. 2014), or antibiotic exposure (Shaw et al. 2019). A fundamental challenge is that community assembly is governed by a confluence of stochastic and deterministic processes. As a result, the structure and function of microbial communities is affected by selection, historical contingency, and chance events, in a manner that remains poorly understood (Costello et al. 2012). Developing a quantitative theory that integrates all of these selective and stochastic ecological processes into an ultimately predictive framework is a major aspiration of microbiome biology. This goal calls for us to identify the quantitative rules governing assembly of microbial communities at different levels of organization.

Recent studies in a range of natural microbiomes, including those of systems as diverse as soils (Nelson, Martiny, and Martiny 2016), the oceans (Louca, Parfrey, and Doebeli 2016), plants (Louca et al. 2016; Burke et al. 2011), and the human gut (Turnbaugh et al. 2009), have compared the metagenomic and taxonomic structure of microbiomes assembled in natural “semi-replicate” habitats (e.g. in different hosts of the same species). These studies have reported intriguing, generic patterns of convergence and variability at different levels of organization. When binned by pathway, the quantitative fraction of the metagenome that is devoted to specific metabolic functions is often found to be quantitatively convergent across habitats, suggesting that these quantitative fractions are the result of environmental selection (Louca et al. 2018). Yet, these studies also find that the taxonomic composition (at the genus or lower level) is highly variable amongst replicate habitats, and this variability was also strong when grouping taxa by their ability to perform specific metabolic pathways (i.e. within functional groups). This has led to the proposal that environmental selection strongly determines the quantitative fractions of the metagenome devoted to different metabolic functions, whereas the taxonomic composition is more variable and sensitive to chance events, environmental heterogeneity, historical contingency, and other sources.

A fundamental limitation of natural surveys is that it is difficult to draw direct mechanistic links between physiological processes at the cellular level and the patterns of convergence and variation at higher levels of organization. We cannot, for instance, explain why the specific ratios of different metabolic pathways are what they are in a given natural environment, nor how they would change in response to specific perturbations. Perhaps the biggest challenge is that, in most natural environments, we simply do not know the exact selective pressures experienced by microbes, nor do we have a detailed chronology of the historical events that may have influenced community assembly.

Thus, a critical step towards quantitatively understanding microbial community assembly at different levels of organization is to study this process in simpler and well-controlled habitats, where all of the selective and non-selective forces at play can be well understood and mechanistic links can be drawn. To this end, we have recently investigated the self-assembly of hundreds of enrichment communities in replicate, identical synthetic habitats of known chemical and physical composition and known assembly history (Goldford et al. 2018). In these experiments, we found a strong convergence across replicates at higher levels of taxonomic organization (i.e. family or higher), despite the presence of substantial variability at lower levels (i.e. genus). In glucose-limited media, for instance, communities adopted quantitatively convergent ratios of the two dominant taxonomic families (Enterobacteriaceae and Pseudomonadaceae) across N~100 replicate habitats, despite the different starting pools of species (inocula) used to colonize each habitat (**Fig. 1A**). At the same time, the species-level composition within each of these families was highly variable, and it was divergent even when communities were started from the same inoculum (Goldford et al. 2018).

**Figure 1.**
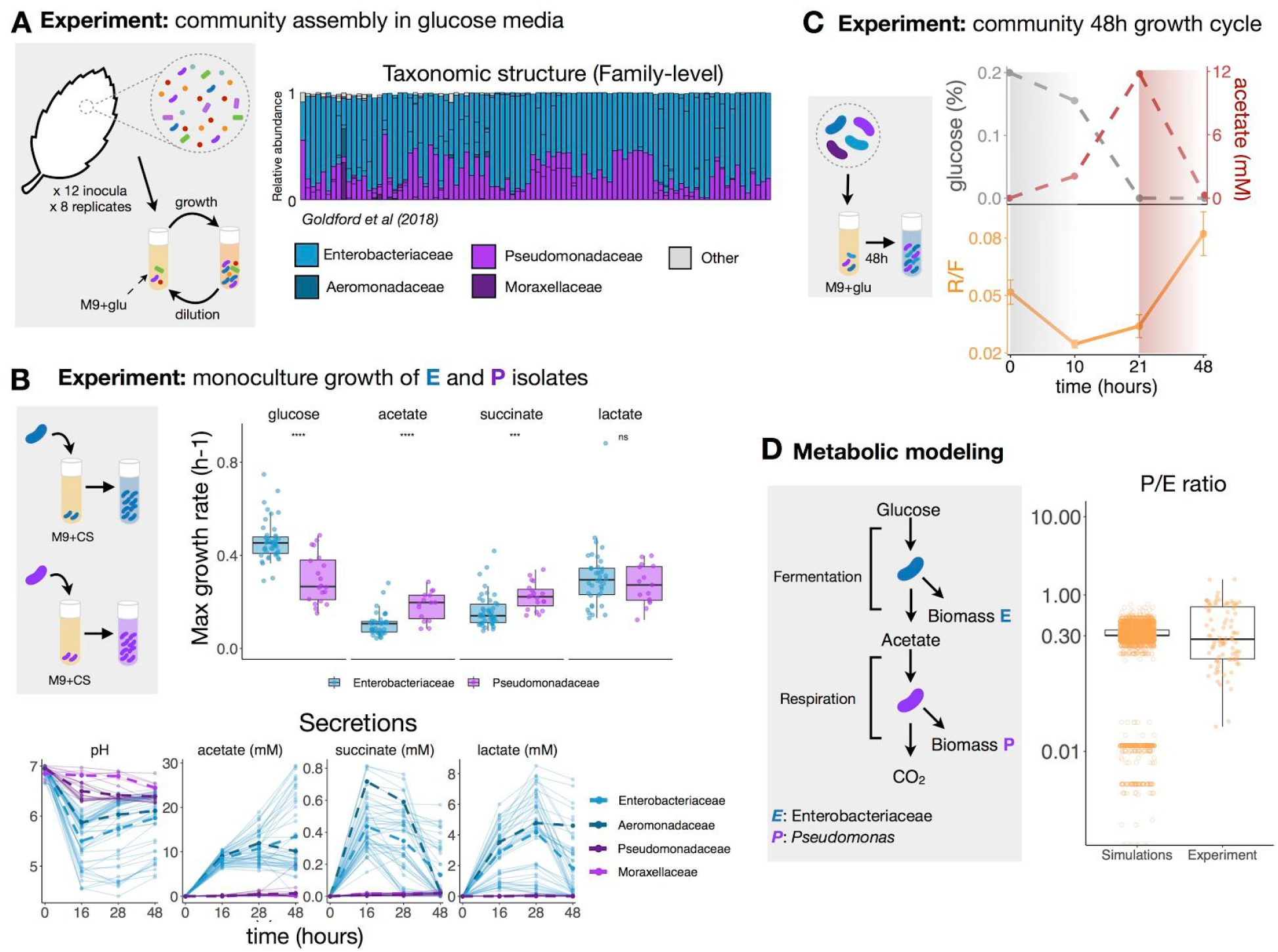
Emergent metabolic structure in self-assembled microbial communities. **A.** Barplots show the relative abundance of the dominant families (Enterobacteriaceae, Pseudomonadaceae, Aeromonadaceae and Moraxellaceae) in 92 communities started from 12 leaf or soil inocula (8 replicates each) after assembly in minimal media with glucose for 12 growth/dilution cycles (data from (Goldford et al. 2018)). Only taxa with relative abundance > 0.01 are shown. Other families are shown in gray. **B.** Isolates belonging to different families were grown in monoculture for 48h in minimal media supplemented with a single carbon source (CS) (glucose, acetate, lactate, or succinate) (N=73, **Fig. S2**). Each dot corresponds to a strain’s maximum growth rate. Note that **** indicates p<=0.0001, *** indicates p<=0.001, ns indicates p>0.05, paired Student’s t-test. We quantified the amount of acetate, lactate and succinate in the medium at various time points for all isolates. The dashed lines represent the mean concentrations for isolates of each family. **C.** Communities were thawed and grown in minimal media with glucose for a single incubation time. Samples were taken at 10h, 21h and 48h, and we measured the R/F ratio and the concentrations of glucose and acetate in the medium. Only one representative community (out of N=9) is shown. See **Fig S7** for other communities. The R/F ratio represents the mean ± sd of the CFU ratios calculated by bootstrapping (N=1000 replicates). **D.** Predicted and observed P/E ratio. *Simulations*: Using Flux Balance Analysis, we calculated the biomass obtained from glucose fermentation by Enterobacteriaceae strains (E) and the biomass obtained from consumption of the *E*’s metabolic byproduct, acetate, by *Pseudomonas* strains (P). The predicted ratio between P and E biomass was calculated for 59 Enterobacteriaceae and 74 *Pseudomonas* metabolic models. The simulations predict a median P/E ratio of ~0.303 (Q1=0.302, Q3=0.356). *Experiment*.: P/E ratio observed experimentally in the glucose communities described in **Fig. 1A** (median=0.27, Q1=0.15, Q3=0.70, N=92). Each dot represents a P/E pair.

These findings, placed in the context of the previously discussed surveys of natural communities, open many questions. What do these family-level assembly rules reflect? Do the convergent ratios of Pseudomonadaceae to Enterobacteriaceae in replicate sugar environments reflect an emergent metabolic self-organization of microbial communities? If so, can we mechanistically explain and quantitatively predict these ratios from first principles? Likewise, why is there such variability at the sub-family level in replicate habitats? Does it reflect neutral drift of species within the same functional guild, or simply random species sampling during colonization, or alternative stable states? (Aguirre de Cárcer 2019; Ley, Peterson, and Gordon 2006; Costello et al. 2012; Fukami 2015; Vellend 2010). Through a combination of new experiments and genome-scale metabolic modeling, we will proceed to address these questions.

## Results

### Emergent metabolic self-organization of microbial community assembly in glucose-limited environments

As previewed above, we have recently found that natural bacterial communities that were serially passaged every 48hr in glucose minimal media for 84 generations (12 transfers) self-assembled into stable communities containing N=2-17 taxa, which coexist thanks to extensive cross-feeding interactions (Goldford et al. 2018). Despite their different starting inocula, these enrichment communities assembled into highly reproducible compositions at the family (or higher) level of taxonomy, while varying widely in their composition at (or below) the genus level (Goldford et al. 2018). Communities were dominated by the Enterobacteriaceae (E) and Pseudomonadaceae (P) families, at a median ratio of P/E=0.27 (N=92, Q1=0.15, Q3=0.70) (**Fig. 1A**).

The reasons for the strong reproducibility of community assembly at higher levels of taxonomic organization, and for the specific quantitative ratios of these two specific families remain unknown. Previous studies in natural communities have found that the fraction of the metagenome engaged in different metabolic functions is also strongly quantitatively conserved among similar habitats (Tringe et al. 2005; Louca et al. 2016, 2018). We thus speculated that the observed taxonomic convergence may reflect an emergent metabolic organization that maps to phylogenetic community assembly through the family-level conservation of quantitative metabolic traits (Aguirre de Cárcer 2019).

To evaluate this possibility, we isolated 73 member strains from 17 of our communities, spanning 11 genera of the families Pseudomonadaceae (N=20 isolates), Enterobacteriaceae (N=47 isolates) as well as other less abundant members such as Moraxellaceae (N=3), Aeromonadaceae (N=1), Alcaligenaceae (N=1), and Comamonadaceae (N=1). On average, our isolates represented 88.8% of the taxonomic composition of the 17 communities from where they were collected (**Fig. S1**; Methods). We first measured the growth rates of all of these isolates in the same glucose M9 minimal medium where the communities had been originally assembled (Methods). All of the 73 isolates were able to grow on glucose in monoculture. Since this is the sole supplied resource, one might conclude that all species are part of the same functional guild of glucose metabolizing bacteria. However, we also found that Enterobacteriaceae have much stronger growth rates than Pseudomonadaceae in glucose medium (median(E, glu)=0.45/h and median(P, glu)=0.27/h, p< 0.0001, **Fig. 1B**, **Fig. S2**). Despite this almost 2-fold difference in growth rate, both families consistently coexist in our synthetic glucose-limited habitats.

In previous work, we found that cross-feeding was key for the maintenance of diversity in our communities (Goldford et al. 2018; Lu et al. 2018). We thus hypothesized that the Pseudomonadaceae may be sustained in the community not because of their ability to metabolize glucose, but rather by their higher competitive ability in the metabolic byproducts secreted by the Enterobacteriaceae. To identify what those byproducts are, we used liquid-chromatography mass spectrometry (LC-MS) to analyze the secreted metabolic byproducts of glucose metabolism for a dominant *Enterobacter* strain in our communities, as well as for *E. coli* MG1655 (Methods). These two species are representative of the two main forms of fermentation typically found in the Enterobacteriaceae (Vivijs et al. 2015). In both cases, we found that acetate, lactate, and succinate were the three primary byproducts secreted into the environment during the exponential phase, consistent with the known patterns of overflow metabolism in Enterobacteriaceae (Vivijs et al. 2015) (**Fig. S3**). Of these, acetate was by far the dominant, at a concentration of 4.7 ± 0.5 mM for *E. coli* and 6.0 ± 0.2 mM for *Enterobacter* after 28h of growth. To test the generality of these secretion patterns, we then proceeded to quantify the amount of acetate, succinate and lactate secreted by the Enterobacteriaceae strains when growing in glucose M9 media (Methods). All three organic acids are strongly secreted by all the Enterobacteriaceae (at similar amounts across all isolates, with some genus-level variation (**Fig. S4**)) but, as expected, not by the Pseudomonadaceae, since they are non-fermentative (**Fig. 1B**). Acetate is in all cases the dominant overflow byproduct (median=8.5 mM, Q1=7.5 mM, and Q3=9.6 mM after 16h of growth).

If the hypothesis outlined above is correct, then Pseudomonadaceae should have a higher growth rate than Enterobacteriaceae in acetate and possibly also in the other organic acids. To test this, we measured the growth rates of all of our isolates in acetate, succinate, and lactate minimal media, respectively (Methods). Pseudomonadaceae did indeed have close to 2-fold median higher growth rates in acetate (median(P, acetate)= 0.20/h and median(E, acetate)= 0.11/h, p<0.0001, paired Student’s t-test), which is the dominant byproduct, and also grew faster in succinate (median(P, succinate)=0.22/h and median(E, succinate)= 0.14/h, p<0.001, paired Student’s t-test), **Fig. 1B**, **Fig. S2** and **Fig. S5**). No significant difference in growth rates in lactate was observed.

Altogether these results suggest that although the Enterobacteriaceae and Pseudomonadaceae found in our communities are all capable of metabolizing glucose, they are not all part of the same “glucose-metabolizers” functional guild. Rather, our findings suggest that Enterobacteriaceae are selected by the glucose added to the medium, due to their fast growth in it. This fast growth is accomplished through an overflow metabolism strategy, which leads to the excretion of highly similar quantities of the same dominant organic acids. This metabolic strategy is conserved at the family level and shared by all of the Enterobacteriaceae in our communities. Finally, our results suggest that the convergent set of organic acids secreted by the Enterobacteriaceae, as a byproduct of their overflow metabolism, provide the main substrate for the growth of Pseudomonadaceae, which are competitively selected by them. Based on this, we propose that the Enterobacteriaceae in our communities (as well as the closely related Aeromonadaceae, which behave in the same way) form a phylogenetically conserved fermenter (F) functional guild, which is selected by the glucose due to their fast growth rates in glucose. In turn, the Pseudomonadaceae (together with the Comamonadaceae and Moraxellaceae) form a second functional guild, which is made up of the respirator (R) bacteria that are selected by the organic acids released by the fermenters, on which they specialize. The amount of acetate excreted by the Enterobacteriaceae species correlates with their maximum growth rate in glucose (**Fig. S6**, R^2^=0.49), further suggesting that selection for the fastest growers in the inoculum inevitably leads to the accumulation of organic acids. It is thus not the ability to metabolize a substrate, but the competitive ability on it, which delineates membership in a functional group.

To further confirm this scenario, we thawed 9 stable communities that had previously been frozen at the end of 12 transfers in the prior study (Goldford et al. 2018). To relieve any potential effects of freezing and thawing on the composition and stability of these communities, we passaged them for an additional three transfers (Methods). We then measured the ratio between respirator (R) and fermenter (F) abundances (the R/F ratio) at different time points during a final 48h growth cycle (at 0, 10, 21, and 48 h), quantifying also the concentrations of glucose and acetate at each time. Consistent with our hypothesis, we find that fermenters have a growth advantage in all communities (characterized by a drop in the R/F ratio) early on the incubation period (T=0-10h) when glucose is abundant (**Fig. 1C**, **Fig. S7**). In turn, respirators have a growth advantage compared to fermenters (characterized by an increase in the R/F ratio) in the second part of the incubation period (T=21h-48h) for most of the communities (7 out of 9) (**Fig. 1C**, **Fig. S7**), when glucose is absent but organic acids are abundant. The other two communities nevertheless exhibit an increase in their R/F ratio from T=10h-T=48h, after glucose had been partially depleted. The growth of the fermenters was accompanied by a depletion of glucose, whereas the growth of the respirators is concomitant with a depletion of acetate (**Fig. 1C**, **Fig. S7**).

Our results indicate that the convergent ratio of Pseudomonadaceae to Enterobacteriaceae (the P/E ratio) reflects the relative frequencies of two metabolic specialist groups: respirators and fermentors (i.e. the R/F ratio). Further supporting this point, in other experiments we have run in our lab in recent years, we have found that when communities lacked either Enterobacteriaceae or Pseudomonadaceae, these families were replaced in highly similar frequencies by members of other families with similar functional roles. For instance, Enterobacteriaceae can be replaced by Aeromonadaceae (**Fig. 1A**, **Fig. S8**), another family of known respiro-fermentative bacteria, which grows strongly in glucose (**Fig. S2**) by fermenting it to the same organic acids as Enterobacteriaceae (**Fig. 1B**). Likewise, in other communities, we have observed that Pseudomonadaceae could be replaced by either Moraxellaceae (**Fig. 1**, **Fig. S8**) or Alcaligenaceae (**Fig. S8**). All of these taxonomically divergent communities have different family compositions, but highly similar convergent ratios of organic acid respirators to glucose fermenters as the one found in **Fig 1A** (R/F=0.29, Q1=0.17, Q3=0.69). This is consistent with the idea that family-level convergence is a proxy for convergent functional self-organization, which arises due to the evolutionary conservation of quantitative metabolic traits such as niche construction, and the growth-rate response to nutrients.

### Genome-scale metabolic modeling quantitatively explains the ratio of both functional groups

This does not explain however why the observed ratio is R/F= 0.29. To test whether this ratio could be explained from first principles, we carried out Flux Balance Analysis simulations using a recently developed extension that naturally exhibits overflow metabolism (Mori et al. 2016). In our simulations, we first used a well-curated genome-scale metabolic model of a representative Enterobacteriaceae, the *E. coli* model (iJO1366, (Orth et al. 2011)), and a well-curated genome-scale metabolic model of Pseudomonadaceae, the *P. putida* model (iJN1463, (Nogales et al. 2020)). Using these models, we determined the biomass ratio of Pseudomonadaceae to Enterobacteriaceae in the limit scenario where E. coli metabolizes all of the glucose, secreting acetate as a byproduct, and P. putida fully respires the produced acetate to CO_2_. The model predicts a *P. putida*/ *E. coli* ratio of P/E = 0.36. Parametric stability analysis shows that this estimate is robust to the specific assumptions made in the model (Methods, **Table S1**). To test how the P/E ratio would vary with different Enterobacteriaceae or *Pseudomonas* species, we compiled a library of 59 Enterobacteriaceae and 74 *Pseudomonas* metabolic models (Methods, and Supplementary Methods) and repeated the above simulation to predict the P/E ratio for every pair of Enterobacteriaceae and *Pseudomonas* (**Fig. S9**). Our simulations predict a median P/E ratio of 0.303 (Q1=0.302, Q3=0.356) which is strongly aligned with the experimentally observed median P/E ratio of 0.27 (Q1=0.15, Q3=0.70, N=92) in our glucose communities (**Fig. 1D**).

### Replaying the tape of community assembly: Non-modular taxonomic variability within functional groups

Despite their quantitatively convergent metabolic self-organization, communities often exhibited substantial taxonomic variation at the genus level or lower, even when they were started from the same inoculum (Goldford et al. 2018). What are the causes of the observed taxonomic variability within each functional metabolic group? One possibility is that the taxonomic variability reflects random sorting of species into different replicate habitats: Some genera may be sampled only into some but not all of the habitats. Since there is no further migration after the first inoculation, this could easily lead to variability, as was already shown elsewhere using simulations (Goldford et al. 2018). An alternative hypothesis is that individual species within the same functional group are “equivalent” and would all have strongly similar fitness and thus exhibit neutral dynamics (Aguirre de Cárcer 2019). Finally, taxonomic variability could also arise due to dynamic multistability in population dynamics, and the existence of alternative stable states (Fukami 2015).

The number of replicates in previous experiments is not high enough to unambiguously discriminate among these alternative hypotheses. Thus, we started a new experiment with 92 replicate communities, all initiated from the same inoculum and propagated in minimal media with glucose as before (**Fig. 2A**). After 18 serial dilution transfers, most communities (77 out of 92) (we will leave the remaining ones aside for now, and return back to them in later sections of this paper) assembled into the metabolic structure described above, consisting of fermenters (glucose specialists) and respirators (organic acid specialists) at similar proportions as we had seen previously (R/F= 0.46, Q1=0.34, Q3=0.65) (**Fig. 2B**). For these communities, we identified two alternative taxonomic compositions within the fermenter functional group and three alternative taxonomic compositions within the respirator functional group (**Fig. 2B**), which are evident by visual inspection and are generally consistent with those found using cluster analysis (**Fig. S10**). The dominant ESVs found in the respirator groups are an ESV of the genus *Alcaligenes* and an ESV of the genus *Pseudomonas*. Among the fermenters, the dominant taxa were two strains of the genus *Klebsiella* (hereafter referred to as *Kp* and *Km* (Methods)).

**Figure 2.**
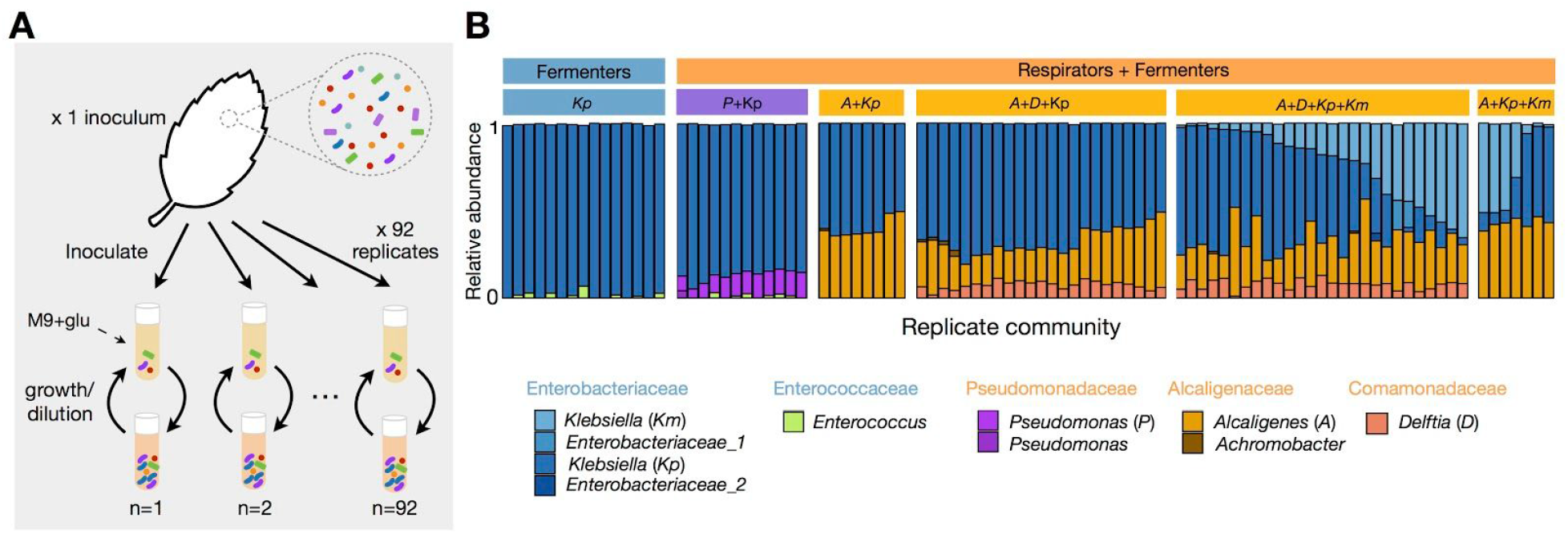
Multiple alternative states at the metabolic and taxonomic level arise from assembly of replicate communities from a single inoculum. **A.** Schematic of experimental design: starting from a highly diverse soil microbial community, 92 communities were serially passaged in replicate habitats with glucose as the single carbon source for 18 incubation (growth/dilution) cycles (48h each). **B.** Taxonomic profile of communities shown at the exact sequence variant (ESV) level (one color per ESV) with corresponding genus and family level assignments. Only the ESVs with a relative abundance > 0.01 are shown. After 18 transfers, we find that replicate communities self-assembled in two major functional groups, fermenters only (N=15) or fermenters with respirators (N=77). Within the fermenter functional group, we can see two alternative taxonomic compositions depending on whether one or two *Klebsiella* strains are present. Within the respirator functional group, we can clearly identify three alternative taxonomic groups (*Pseudomonas*, *Alcaligenes*, and *Alcaligenes* + *Delftia*).

Importantly, the taxonomic compositions of the fermenter and respirator groups in a given community were not independent (p_obs>null_ <0.001 using C-score, Methods, **Fig. S11**), indicating that community assembly is not modular. In communities where *Alcaligenes* dominated the respirator group, the fermenter group could contain either *Kp* alone or both *Kp* and *Km* above a 0.01 abundance threshold. By contrast, when *Pseudomonas* dominates the respirator group, the fermenter taxa would only contain one of those strains (*Kp*), but never the other, and it also may contain a strain of the genus *Enterococcus*, which is in turn never found co-occurring with *Alcaligenes*. The composition of the respirator group is also strongly determined by its dominant taxa: *Alcaligenes* may co-occur with *Delftia* (in 50 communities), *Achromobacter* (N=7), and often both (N=6). *Pseudomonas*, on the other hand, is never found together with neither of them. Importantly, *Alcaligenes* and *Pseudomonas* were never found together above an abundance of 0.01.

### Mutual exclusion between two respirator strains drives alternative states in multispecies communities

Why do some communities contain *Alcaligenes* as the main organic acid specialist while others contain *Pseudomonas*? A first hypothesis is that these alternative states arise due to random sampling from the initial species pool. Because there is no immigration, any community where *Alcaligenes* and/or *Pseudomonas* was not stochastically sampled during inoculation will inevitably not have those strains at at the end of the experiment (Goldford et al. 2018). To test this hypothesis, we first ask if, consistent with this hypothesis, communities dominated by *Alcaligenes* have no *Pseudomonas*, and similarly, if communities dominated by *Pseudomonas* have no *Alcaligenes*. Contrary to this hypothesis, we find that in ~28% (18/65) of the *Alcaligenes* dominated communities, *Pseudomonas* is present below an abundance of 0.01 but above a cutoff that corrects for spurious detection during amplicon sequencing (Methods, **Fig. S12**). Likewise, in ~67% (8/12) of the *Pseudomonas* dominated communities, *Alcaligenes* is present below an abundance of 0.01, and above the error threshold (Methods, **Fig. S12**). This result suggests that the alternative states we observed are generally not caused by random sampling, nor to the extinction of established taxa due to the population bottleneck applied by every serial dilution into fresh media.

The existence of a state where *Alcaligenes* is abundant and *Pseudomonas* is rare and of a state where *Pseudomonas* is abundant and *Alcaligenes* is rare, in addition to the lack of a state where *Alcaligenes* and *Pseudomonas* are both abundant, suggests that *Alcaligenes* and *Pseudomonas* may mutually outcompete one another, which can give rise to multistability in population dynamics. This can also lead to alternative stable states within a functional group (Gonze et al. 2017; Fukami 2015; Case 1990; Shaw et al. 2019; Chen et al. 2014). To test the possibility of multistability, we isolated the dominant strains (*Kp*, *P*, and *A*) and inoculated multiple populations of *Klebsiella* (*Kp*) with varying initial densities of *Alcaligenes* and/or *Pseudomonas* (**Fig. 3A**), mapping a two-dimensional phase portrait (**Fig. 3B** and **Fig. S13-S14**). We then passaged them in minimal glucose media for 12 growth-dilution cycles, allowing the communities to reach an equilibrium, and measured their abundances at three timepoints by colony counting (**Fig. 3A**) (Methods).

**Figure 3.**
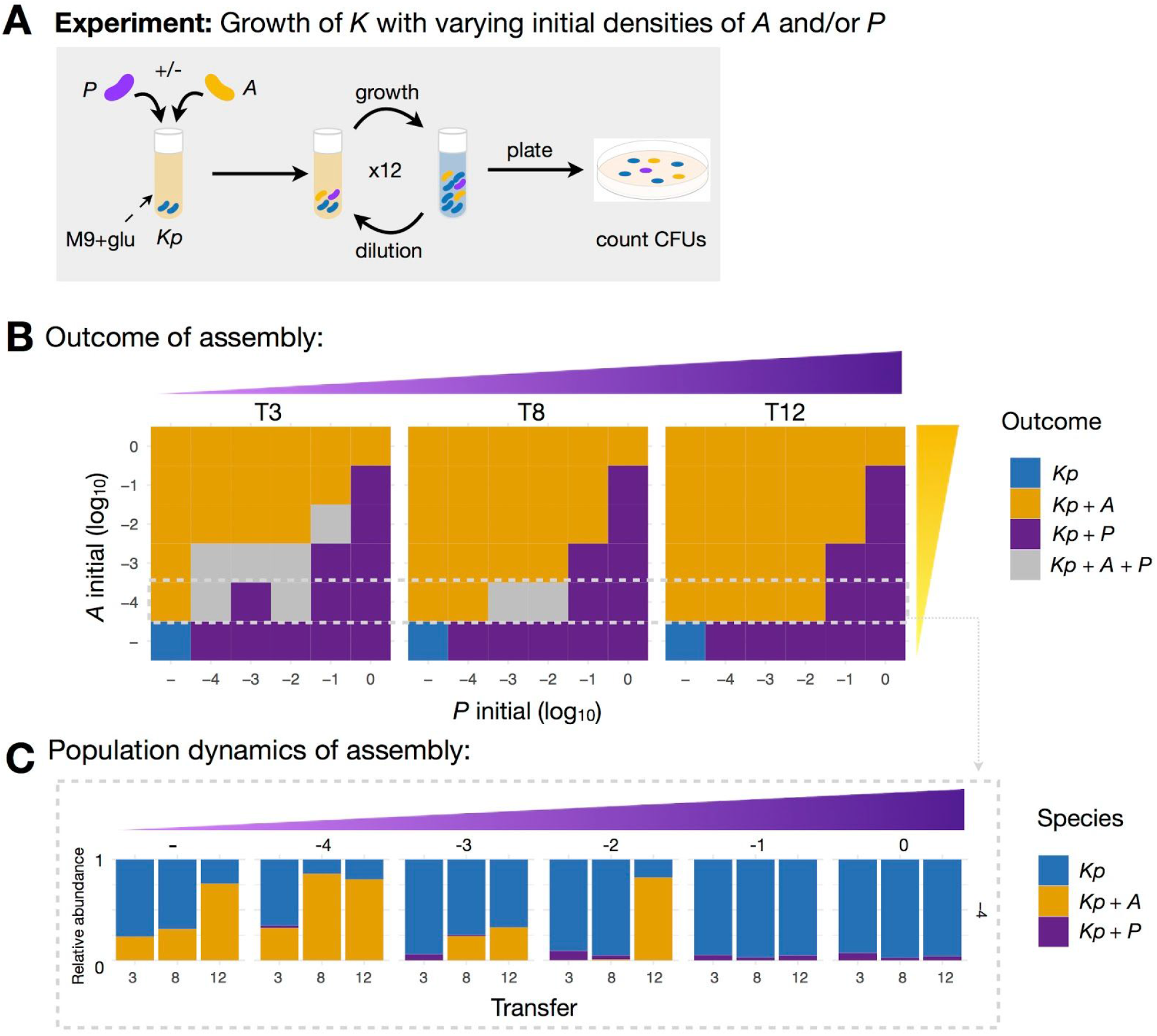
Multistability leads to alternative attractors in community composition. **A.** We isolated the three dominant strains - *Klebsiella* (*Kp*), *Alcaligenes* (*A*), and *Pseudomonas* (*P*) that make up the two major alternative attractors, and grew them in pairwise coculture (*Kp*+*A* or *Kp*+*P*) or in three species consortia (*Kp*+*A*+*P*) by mixing *Kp* with different initial densities of *A* and/or *P* (see Methods). These reconstituted communities were grown in the same conditions as the top-down assembly communities for 12 transfers (Methods). **B.** Phase portrait showing the state of the community after T= 3, 8, and 12 transfers. A square is colored orange if a community that was started there contained *A* but not *P* at time T. It is purple if it contained *P* but not *A*, and it is gray if both *A* and *P* were present. We can see that the phase portrait is divided in two regions: The upper-left diagonal is made up by the basin of attraction of *Alcaligenes* dominated communities, whereas the bottom-right diagonal contains the basin of attraction for *Pseudomonas* dominated communities. *Alcaligenes* and *Pseudomonas* mutually exclude each other depending on their starting densities. See **Fig. S13** for a second biological replicate experiment, which gives similar outcomes. **C.** Temporal dynamics of the relative abundance of each taxa for a subset of the communities shown in **B**. See **Fig. S14** for the time series of all pairwise initial conditions of the phase portrait.

Consistent with the expectation of multistable population dynamics, we find that *Pseudomonas* and *Alcaligenes* can both coexist with *Klebsiella* individually but not together, regardless of their initial densities (**Fig. 3B-C**, and **Figs S13 and S14**). Importantly, and as expected from the multistability hypothesis, the invasion outcome (i.e. whether *Pseudomonas* or *Alcaligenes* dominate) depends on the initial position of the population in the phase portrait (i.e. the relative abundances of *Pseudomonas* and *Alcaligenes*). In the upper left part of the portrait when *Pseudomonas* starts at low and *Alcaligenes* at high abundance, communities converge to a state dominated by *Alcaligenes*. In the lower-right part of the phase portrait, communities converge to a state dominated by *Pseudomonas* (**Fig. 3B**).

In a multistable system, a switch from one state to another only happens in the presence of strong perturbations, for instance, by changing the abiotic environment (e.g., diet shift) or by changing the biotic environment (e.g. immigration of new species, or extinction of established species). Given that our communities were assembled under highly-controlled chemo-physical conditions without migration, we expect that once a community has switched to one state it remains in that state, and thus switches from the high *Alcaligenes*/low *Pseudomonas* to the low *Alcaligenes*/high *Pseudomonas* state, or vice-versa, should be rare. To test this in our communities, we sequenced the full temporal dynamics for a representative subset of the communities (**Fig. S15**). The dynamics reveal that *Alcaligenes* either blooms early during the assembly of the community, after which it remains at high relative abundance (mean=0.27 ± 0.11), or it never invades and remains at low abundance (mean=0.0027 ± 0.0019), in which case *Pseudomonas* may invade and take the role of the dominant organic acid specialist (mean= 0.12 ± 0.030). Furthermore, the location of the basins of attraction for *Alcaligenes* and for *Pseudomonas* inferred from the phase portrait obtained with the bottom-up reconstitution of a minimal community (*Kp*, *A*, *P*) (yellow and purple regions, respectively, in **Fig 3**) approximates well the temporal community trajectories (**Fig. 4**). Communities generally start in the transition region, which represents the unstable region between both stable states. As communities self-assemble, they approach one of the attractors and never switch between states (**Fig. 4B**). Together, these findings suggest that the alternative states we observed between the two organic acid specialists likely arise from dynamic multistability driven by mutual inhibition between them. Whether *Alcaligenes* or *Pseudomonas* invades seems to be stochastic, but once invasion occurs (‘tipping point’), the community state is maintained over time.

**Figure 4.**
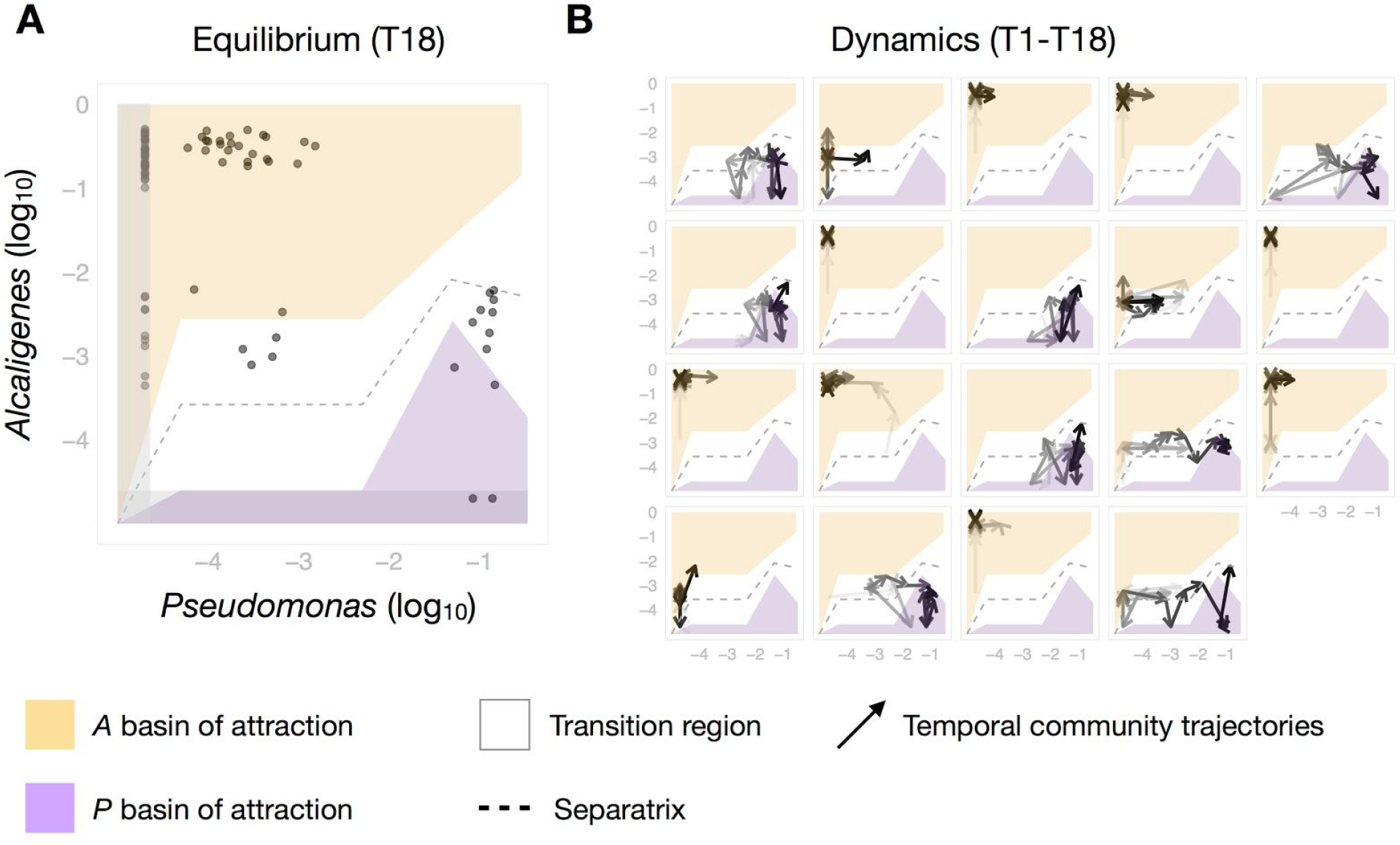
Multistable metabolic attractors between two organic acid specialists. Phase diagram inferred from the bottom-up experiment in **Fig. 3** (Methods), showing the basins of attraction for *Alcaligenes* dominated (yellow area) and for *Pseudomonas* dominated (purple area) states, separated by a transition region (white area). The gray dashed line indicates the separatrix between the two basins of attraction. In **A**, the dots show the relative abundance of *Alcaligenes* and *Pseudomonas* at Transfer 18 (n=92). The gray shaded areas indicate the regions of low Alcaligenes and low Pseudomonas that are below the detection level of amplicon sequencing. In **B**, overlaid are the trajectories of the relative abundance of *Alcaligenes* and *Pseudomonas* for all communities for which we measured time series population dynamics (N=19). The arrows become darker with time (i.e. from T1 to T18).

### The presence of alternative respirators affects interactions among members of the fermenter group

As discussed above, the identity of the dominant member of the respirator group shapes the composition of the fermenter group, indicating that the assembly of both taxonomic groups within a community is not modular (not independent from each other) (**Fig. S11**). Communities with *Pseudomonas* as the dominant respirator co-occur with a single ESV of *Klebsiella* (*Kp*) as the sole member of the fermenter group, whereas when *Alcaligenes* dominates it can co-occur with both *Kp* and *Km*. This suggests that Alcaligenes and Pseudomonas may (differently) modulate interactions between *Kp* and *Km*, a signature of higher-order interactions (Sanchez-Gorostiaga et al. 2019; Billick and Case 1994; Guo and Boedicker 2016; Bairey, Kelsic, and Kishony 2016; Sanchez 2019; Mickalide and Kuehn 2019; Letten and Stouffer 2019; Tekin, Yeh, and Savage 2018). To test this possibility, we grew *Kp* and *Km* together in pairwise coculture, as well as in three species consortia together with either *Alcaligenes* or with *Pseudomonas*. Communities were passaged every 48h for 6 transfers in minimal glucose media (Methods). We found that *Km* is competitively excluded by *Kp* in pairwise co-culture and, consistent with the patterns we found in self-assembled communities, the species exclude each other when *Pseudomonas* was present (**Fig. S16**). However, in the presence of *Alcaligenes*, the three strains can coexist. This suggests that *Alcaligenes* buffers (i.e. neutralizes) the competition between the two *Klebsiella* strains (**Fig. S16**).

### Migration between communities leads to strong metabolic convergence

A non-negligible number of the 92 replicate self-assembled communities (15/92) shown in **Fig. 2B** assembled into a functional state where respirators comprised less than 1% of the population (R/F=0.002, Q1=0.0014, Q3=0.0042). The existence of these communities appears to be a violation of the metabolic assembly rule discussed in **Fig. 1**. It is not clear, however, whether such communities represent a stable metabolic attractor, as opposed to being “frozen” in a transient or meta-stable state as a result of community assembly in a closed system. Previous theoretical and experimental work has shown that dispersal between communities can homogenize communities, and disfavor marginally stable equilibria (eg. (Chase 2003; Fodelianakis et al. 2019; Leibold et al. 2004; Stegen et al. 2013)). Therefore, we hypothesized that opening the system by connecting communities through migration may prevent the system from getting locked into a meta-stable state, recovering the metabolic assembly rule.

To investigate this hypothesis, we repeated our community assembly experiment using the same initial inoculum, but this time, in addition to the normal transfer, we also imposed migration between communities for twelve growth cycles and then allowed the communities to stabilize without migration for six additional transfers (**Fig. 5A**). This is similar to a metacommunity model (Leibold et al. 2004), and hereafter referred to as ‘global migration’. As predicted, we found that after 12 growth cycles of assembly under constant global migration, communities strongly converged to a single metabolic attractor composed of fermenters and respirators (**Fig. 5B**). At the taxonomic level, they converge again to the most common state observed in the ‘no migration’ case (**Fig. 2B**), that is, the state dominated by the Enterobacteriaceae and Alcaligenaceae families and a median R/F ratio of 0.41 (Q1=0.37 and Q3=0.44) (**Fig. 5B**). The community composition and R/F ratio remain quantitatively similar after six transfers of stabilization without migration (R/F=0.40, Q1=0.37, and Q3=0.47) despite a timid increase in the abundance of *Delftia* (~0.04) (**Fig. 5B**).

**Figure 5.**
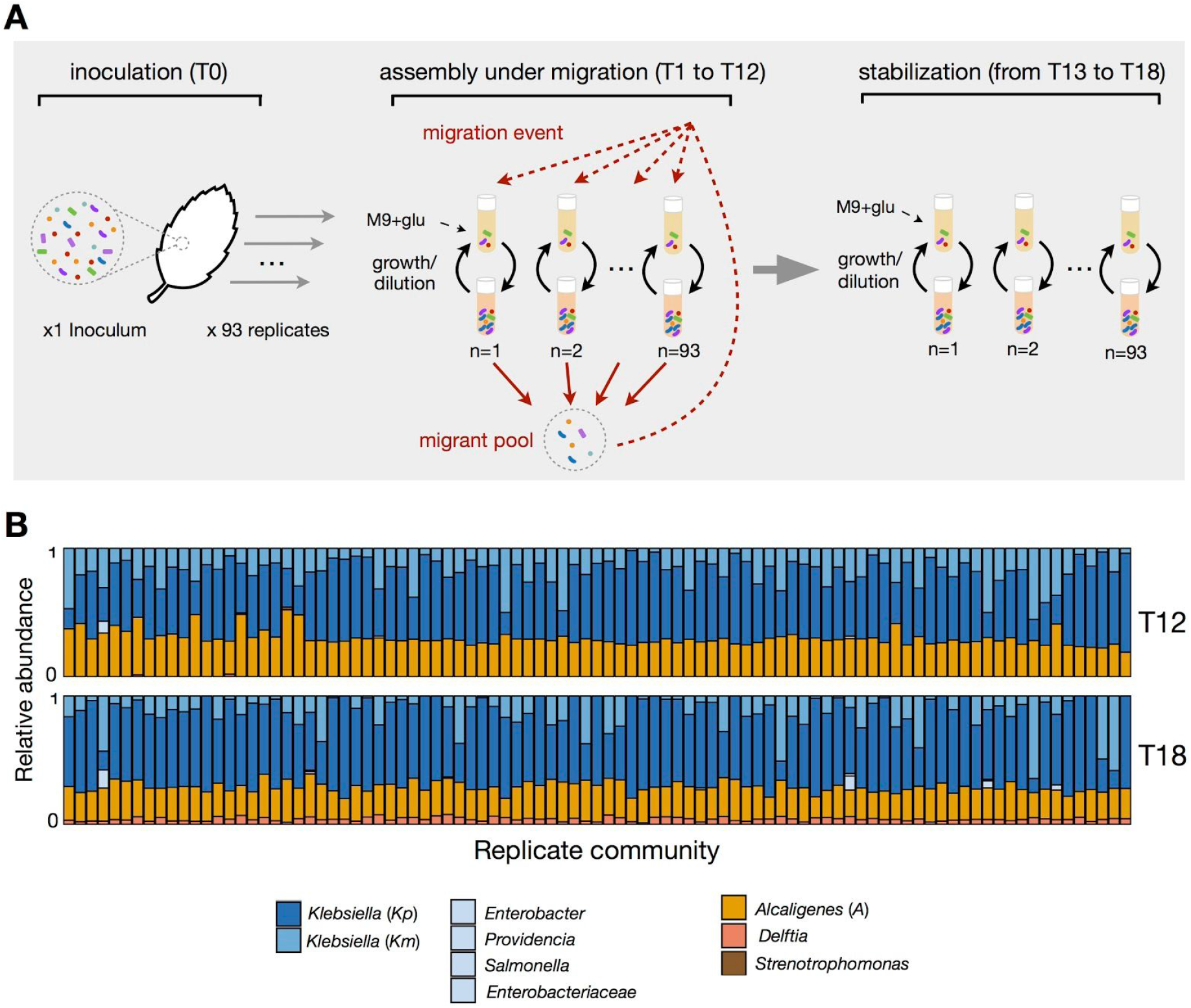
Migration between communities leads to strong metabolic and taxonomic convergence. **A.** 93 replicate communities all started from the same inoculum (and the same inoculum as in **Fig. 2**) were assembled in an open system with global migration - that is, in addition to the normal transfer, each community received a small amount of migrants from a common migrant pool. Communities were assembled under this migration scenario for twelve growth cycles (T1-T12), after which migration was stopped and communities were allowed to stabilize for six additional transfers without migration (T13-T18). **B.** Taxonomic composition of the communities at Transfers 12 and 18. Only the ESVs with a relative abundance > 0.01 are shown.

This result is consistent with the hypothesis that the low R/F (~0.002) functional state observed in this experiment is dynamically metastable, and that communities would converge to the same emergent metabolic attractor as those in previous experiments, given enough time. We again find that both *Klebsiella* (*Kp* and *Km*) and *Delftia* can co-occur with *Alcaligenes*, providing further evidence to the idea that alternative stable states at the species level can arise from mutual exclusion among members of the subdominant, respirator functional group, which in turn recruit different sets of additional taxa in both the respirator and fermenter groups.

## Discussion

In this paper, we set out to mechanistically investigate the patterns of convergence and divergence observed at different levels of microbial community organization. Multiple surveys of natural environments had previously found a pattern of emergent simplicity in microbial community assembly, whereby the microbiome seems highly variable at the species level, but becomes predictable at higher levels of organization, e.g. when we bin the metagenome by metabolic function (Louca, Parfrey, and Doebeli 2016; Louca et al. 2018). The complexity and poor understanding of selective pressures and assembly histories in natural habitats has made it difficult to mechanistically and quantitatively understand these patterns. For instance, the specific ratios of various functional groups in a given habitat could not be explained from first principles, nor reconciled with known cellular processes and biochemical mechanisms. Here, we have addressed this challenge by focusing on communities assembled on synthetic habitats via serial passaging, where the selective pressures can be determined. We first show that, in glucose-limited synthetic environments, the quantitatively convergent ratio of Pseudomonadaceae to Enterobacteriaceae observed in previous studies reflects an emergent metabolic self-organization between glucose specialists and organic acids specialists. Enterobacteriaceae grow faster on glucose than Pseudomonadaceae. This fast growth in glucose is achieved through a respiro-fermentative overflow metabolism, which leads to the release of high-energy metabolic intermediates (mainly acetate, succinate and lactate) into the environment. These metabolic by-products in turn create new resource niches for Pseudomonadaceae, who generally grow faster on the organic acids than Enterobacteriaceae. Further, we show that the ratio of Pseudomonadaceae/ Enterobacteriaceae observed experimentally is in agreement with the ratio predicted from genome-scale models with zero fitting parameters.

This emergent self-organization between glucose specialists and organic acids specialists after ~60 generations of community assembly in a glucose-limited environment shows a remarkably close parallel with findings from long-term experimental evolution of a single species. A notable example is the evolution of an acetate cross-feeding polymorphism in *E. coli*. Starting with a single strain of *E. coli*, growth in a glucose-limited environment repeatedly leads to the evolution of a glucose specialist (which exhibits both an increased rate of glucose uptake and of acetate excretion), and of an acetate cross-feeder with an enhanced ability to use the acetate released by the glucose specialist (Rosenzweig et al. 1994; Turner, Souza, and Lenski 1996; Treves, Manning, and Adams 1998; Rozen and Lenski 2000). This suggests that the emergent self-organization in top-down community assembly in glucose mirrors the bottom-up emergence of the same metabolic structure by evolutionary forces, further reinforcing the predictability of such structure. Intriguingly, strongly similar quantitative patterns of coexistence between fermenters and respirators, and more specifically, Enterobacteriaceae and Pseudomonadaceae, has also been observed in host-associated microbiomes, such as in the apple stigma microbiome (a sugar-rich environment) (Cui et al. 2020) and the microbiome of *C. elegans* (Berg et al. 2016).

To understand why there is substantial variability at lower levels of taxonomic organization (genus or lower level) within each functional group, we monitored the self-assembly of hundreds of replicate communities that were initially seeded from the same inoculum. Different mechanisms have been proposed to explain the emergence of alternative states under such constant conditions: random sampling from the initial species pool, neutral interactions between functionally redundant species and multistability (Aguirre de Cárcer 2019; Goldford et al. 2018; Costello et al. 2012). We tested these different hypotheses in our glucose communities, and showed that the observed taxonomic variability was not due to random sampling from the initial species pool nor to neutral community dynamics but arose due to dynamical multistability caused by mutual exclusion (Leventhal et al. 2018) between two dominant respirator taxa, which in turn lead to further taxonomic variability within the fermenter metabolic group.

Our results also indicated that the taxonomic compositions of each of the two functional groups in a community are not independent, but rather they can strongly affect one another, indicating that community assembly is not modular in our experiments (Grilli, Rogers, and Allesina 2016; Enke et al. 2019). Interestingly, the main driver of taxonomic variability among replicates was the dominant member of the respirator group (a sub-dominant species). Amelioration of competition between two fermenter strains in the presence of one (but not the other) dominant respirator points to the subtle role that high-order interactions may play in community assembly (Billick and Case 1994; Mickalide and Kuehn 2019; Sanchez 2019; Sanchez-Gorostiaga et al. 2019; Senay et al. 2019; Levine et al. 2017; Grilli et al. 2017). This is also in line with previous observations of the potential importance of sub-dominant bacteria in shaping the composition of microbial communities (Lu et al. 2018).

Finally, our results make a case for the quantitative definition of functional niches: all members of the Enterobacteriaceae and Pseudomonadaceae groups are capable of metabolizing both the supplied sugars and the endogenously produced organic acids. It is not the qualitative difference in their metabolic niches, but rather the strong (~2-fold) differences in fitness between both families in each substrate, what delineates the two different functional groups. Not surprisingly, members of the Pseudomonadaceae family are also known to choose organic acids over sugars when both are available (Bajic and Sanchez 2020; Rojo 2010). The importance of diauxic switching for coexistence, particularly in periodic environments such as those used in our study, is predicted in numerous models but still relatively unexplored experimentally (Pacciani-Mori et al. 2019; Goyal, Dubinkina, and Maslov 2018).

Altogether, our work suggests that the observed convergence at higher levels of community organization reflects the existence of global metabolic attractors in microbial community assembly, which can be quantitatively predicted from first principles of bacterial metabolism. Efforts to formulate a predictive, quantitative theory of microbial community assembly require us to identify the right level of description at which community assembly is reproducible and thus predictable. The confluence between our findings and recent empirical results in natural communities assembled in nutritionally similar habitats (Cui et al. 2020) point to promising directions where in vitro and in vivo work can cross-inform each other and be used together to develop a theory of microbial community assembly.

## Methods

### Isolation of strains

Strains were isolated from several communities previously stabilized in glucose minimal media and stored in 40% glycerol at −80C. The communities used were C1R2, C1R4, C1R6, C1R7, C2R4, C2R6, C2R8, C4R1, C7R1, C8R2, C8R4, C8R5, C10R2, C11R1, C11R2, C11R5, C11R6, where CXRY stands for initial environmental sample (inoculum) X replicate community Y (Goldford et al. 2018). These communities were plated in three different media: Tryptic Soy Agar (TSA) and minimal M9 supplemented with glucose or citrate at concentration 0.07 moles of carbon per liter. Isolates from these plates were streaked on the corresponding medium based on visual inspection of colony morphology after 2 and 5 days. Colonies from the streaked plates were streaked twice more on new plates, then cultured in the corresponding liquid medium (Tryptic Soy Broth (TSB), M9 glucose or M9 citrate) and stored at −80C with 40% glycerol.

### Growth curves and maximum growth rate calculation

Isolates were streaked from glycerol on TSA plates and grown at 30C for 48h. Single colonies of each isolate were used to inoculate 500uL TSB in a deep-well plate. These pre-cultures were incubated at 30C without shaking for 48h. Pre-cultures were then diluted 1:1000 in M9 supplemented with either glucose, acetate, lactate, or succinate at a final concentration of 0.07 moles of carbon per liter. The final volume for the growth assays was 100uL in 384 well plates. Each isolate was assayed in two replicates. For computing the maximal exponential growth rate, each replicate was first smoothed by fitting a generalized additive model with an adaptive smoother, using the *gam* function from the *mgcv* package in R.

### LC-MS of E. coli and Enterobacter on minimal glucose

*E. coli* MG1655 and an Enterobacter strain isolated from the glucose communities in (Goldford et al) were revived from frozen stock by streaking on LB Agar. Two replicate colonies of each strain were used to inoculate separate 50ml falcon tubes which contained 5ml of LB-Lennox and were incubated at 30C overnight (shaking at 200RPM). After ~16h of growth, overnight cultures were brought into balanced exponential by three serial transfers into fresh LB (1ml of culture in 4ml of fresh media). The first two transfers were performed at 1h intervals whilst after the final transfer the cultures were allowed to grow for 1h and 30 min. Cells were centrifuged, washed and re-suspended 3 times, using 1.1× M9 media (containing no carbon source). After the final washing step, cells were normalized to an OD620 of 0.1. 500ul of M9 glucose in a 96 deep well plate was inoculated with 4ul of normalized cells, and grown at 30C. After 28h of growth, spent media was extracted using 0.2um Acropep filter plates. Spent media was submitted for a targeted metabolomics analysis carried out by the Metabolomics Innovation Center (TMIC), in Alberta, Canada. This analysis quantified the abundance of >140 metabolites including biogenic amines, amino acids, acylcarnitines, glycerophospholipids, and organic acids by LC-MS.

### 48h growth assay of single strains

Strains were revived from frozen stock and acclimated to growth on glucose minimal media (500uL) for 48h. 4uL of the grown cultures were then inoculated into 500uL fresh glucose media (4 replicates each) and samples were collected at different times during the 48h growth cycle (one replicate used per timepoint, i.e. at 0h, 16h, 28h, and 48h). At each timepoint, 100uL of samples were collected and stored at −80C with 40% glycerol. The remaining samples were immediately centrifuged at 3000 rpm for 25min to separate the cells from the supernatant. The supernatant was transferred to a 96 well 0.2μm Acroprep filter plate on top of a 96 well NUNC plate fitted with the metal collar adaptor and centrifuged at 3000 rpm for 10 min. The supernatant was immediately frozen at −80C until processing.

### 48h growth assay of communities

Previously stabilized communities in glucose minimal media for 12 serial transfers (Goldford et al. 2018) were revived from frozen stock and serially transferred for three passages on glucose minimal media, under the same experimental conditions as before. We selected a subset (N=9) of communities where fermenters and respirators were detected after three serial transfers. At the start of the fourth passage, 4 replicate 96-wellplates were started. Samples were collected at different times during the 48h growth cycle (one plate used per timepoint, i.e. at 0h, 10h, 21h and 48h). At each timepoint, 100ul of samples were taken and stored at −80C with 40% glycerol. The remaining samples were immediately centrifuged at 3000 rpm for 25min to separate the cells from the supernatant. The supernatant was processed as described above.

### Measurement of glucose, acetate, lactate and succinate concentrations

Glucose concentration was measured using the glucose GO assay kit from Millipore (GAGO20). Acetate concentration was measured using the Acetate assay kit (ab204719). D-lactate concentration was measured using the D-Lactate assay kit (ab83429). Succinate concentration was measured using the Succinate assay kit (ab204718). For each assay, the supernatant was diluted (if needed) to ensure that the OD readings are within the standard curve range.

### pH measurement

Determination of pH was done using the same filtered supernatant as for the assays described above. Bromocresol purple (BCP) was used as a pH indicator. The standard curve was prepared by adding different amounts of HCl 1M to M9 without carbon, and measuring pH with an electronic pH-meter. pH of the samples was interpolated on the standard curve as described in (Yao and Byrne 2001).

### Fermentation profile assignment

We assigned each Family to a fermentation profile-respirator (R) or fermenter (F) (**Table S2**). For instance, bacterial genera belonging to the Enterobacteriaceae family are well-known fermenters while bacterial species belonging to the genus *Pseudomonas* are well-known non-fermenters. When counting CFUs, R and F were distinguished by platting on chromogenic agar (HiCrome Universal differential Medium from Sigma). White colonies were counted as R and blue/purple colonies were counted as F. Each isolated strain was platted on chromogenic agar to confirm its R/F assignment. There is a positive correlation between the R/F ratio obtained by CFU counting and 16S sequencing (slope= 1.51, intercept −0.81, R^2^=0.49; **Fig. S17**).

### FBA and constrained-based modeling

Using genome scale metabolic modeling, we can obtain quantitative predictions of the biomass obtainable from glucose fermentation by an *E. coli* metabolic model (such as iJO1366) as well as the biomass obtainable from consumption of *E. coli* metabolic by-products by a *P. putida* metabolic model (such as iJN1463). To predict the amount of *E. coli* biomass generated, we simulate growth on excess glucose using constrained allocation flux balance analysis (CAFBA) (Mori et al. 2016). We use CAFBA because it includes a proteome-allocation constraint that results in the secretion of organic acids such as acetate in aerobic environments. The CAFBA predicted secretions are then used to set the environment for the *P. putida* model. To predict the amount of *P. putida* biomass we simulate growth on the constructed environment using Flux Balance Analysis (see Supplementary Methods). For the *E. coli* (iJO1366) and *P. putida* (iJN1463) metabolic models we obtain a predicted P/E ratio of 0.36, which is close to the empirically observed median P/E ratio of 0.3 for our glucose communities (**Fig 1A, 1D**, (Goldford et al. 2018)). This ratio is robust to large (2 fold) changes in the parameter used for CAFBA simulations (**Table S1**). To test how the P/E ratio would vary with different Enterobacteriaceae or Pseudomonadaceae species, we compiled a library of 59 Enterobacteriaceae and 74 Pseudomonadaceae metabolic models and predicted the P/E Ratio for every pair of models (**Fig 1D**). The predicted ratios have a median of 0.3 and display a similar bimodal distribution to that observed in our glucose communities. Similar distributions are seen if we separately examine each genus of Enterobacteriaceae, or species of *Pseudomonas* (**Fig S9**).

### Sample collection

A soil sample was collected from a natural site in West Haven (CT, USA) using sterilized spatula, placed into a sterile bottle, and returned to the lab. 10 g of soil sample were then placed into a new sterile bottle and soaked into 100mL of sterile PBS supplemented with 200 μg/mL cycloheximide to inhibit eukaryotic growth. The bottle was vortexed and allowed to sit for 48 hours at room temperature. After 48h, samples of the supernatant solution containing the ‘source’ soil microbiome were used as inoculum for the experiment (see section below) or stored at −80C after mixing with 40% glycerol.

### Growth medium and ‘no migration’ experimental setup

Replicate communities from the same source community were cultured separately in the wells of 96 deep-well plates (VWR). Each replicate community was initiated by inoculating 4uL from the source community into 500uL of M9 minimal media supplemented with glucose 0.2% (i.e., 0.07 C-mol/L) (as in (Goldford et al. 2018)). The communities were grown at 30C under static conditions for 48h. After 48h growth, 4uL from the grown culture was transferred to fresh media. This dilution-growth cycle was repeated 18 times. For the first two growth cycles, cycloheximide (200 μg/mL) was added to the media. OD620 was measured at the end of each growth cycle and samples of the grown communities were stored at −80C after mixing with 40% glycerol.

### Migration between local communities experiment

Similar to the treatment without migration, each replicate community was initiated by inoculating 4uL from the source community into 500uL of M9 minimal media. At the end of each growth cycle, however, 4uL from each well was pooled into a ‘migrant pool community’ and diluted 10000-fold. Each well of the fresh media was inoculated with 4uL of this migrant pool community in addition to the 4uL from the corresponding replicate community (well) from the previous growth cycle. Thus, each replicate community from the 96 deep-well plate represents a local community from the same meta-community where the local communities are linked through migration. This migration step was performed for the first 12 growth cycles, followed by 6 dilution-growth cycles without migration (normal transfer only). OD620 was measured at the end of each growth cycle and culture samples were stored at −80C after mixing with 40% glycerol.

### DNA extraction and library preparation

Samples to be sequenced were centrifuged for 30min at 3500rpm. DNA extraction was performed following the DNeasy 96 Blood & Tissue kit protocol for animal tissues (QIAGEN) including the pre-treatment step for Gram-positive bacteria. DNA concentration was quantified using the Quan-iT PicoGreen dsDNA Assay kit (Molecular Probes, Inc) and normalized to 5ng/uL. 16S rRNA amplicon library preparation was conducted using a dual-index paired-end approach (Kozich et al. 2013) and has been described in detail in (Goldford et al. 2018). The PCR reaction products were purified and normalized using the SequalPrep PCR cleanup and normalization kit (Invitrogen).

### Sequencing and taxonomy assignment

The community composition profile was based on 16S (V4) rRNA gene sequence analysis, a commonly used genetic marker for classifying bacteria as it is highly conserved between different species. The samples were sequenced at the Yale Center for Genome Analysis (YCGA) using the Illumina MiSeq (2×250 bp paired-end) sequencing platform. Post-sequencing processing of the raw sequences, namely demultiplexing and removing the barcodes, indexes and primers, was performed using QIIME (version 1.9, (Caporaso et al. 2010)). The generated fastq files for the forward and reverse sequences were analysed using the Dada2 pipeline (version 1.6.0) to infer exact sequence variants (ESVs) (Callahan et al. 2016). The forward and reverse reads were trimmed at position 240 and 160, respectively, and then merged with a minimum overlap of 100bp. All other parameters were set to the Dada2 default values. Chimeras were removed using the “consensus” method in Dada2. The taxonomy of each 16S exact sequence variant (ESV) was then assigned using the naïve Bayesian classifier method (Wang et al. 2007) and the Silva reference database version 128 (Quast et al. 2013) as described in Dada2. A single strain *E. coli* (n=10) and a commercial DNA mock community (D6305, Zymo Research, Irvin, CA, USA) (n=12) were used as positive controls to correct for spurious detection during amplicon sequencing (**Fig. S12**).

### Species co-occurrence analysis

Null model analysis of species co-occurrence for our assembled communities were carried out using the function *cooc*_*null*_*model* of the *EcoSim* R package (Gotelli 2000) with metric set to C-score. We first constructed a presence-absence matrix with all dominant genera (relative abundance >0.01) as rows (except for *Kp* and *Km* that were considered as two separate genera) and all replicate communities (N=92) as columns. We compared the observed C-score for our communities with the C-scores generated from 1000 randomly constructed null assemblages (matrices) using the fixed-equiprobable algorithm (*Sim 2*).

### Isolation of dominant taxa

We isolated the four most abundant ESVs, two belonging to the Enterobacteriaceae family, one *Pseudomonas* and one *Alcaligenes*, and verified their taxonomy by sequencing the full-length 16S rRNA gene (GENEWIZ). Taxonomy was assigned using both the Silva database (v1.2.11) and the RDP Naive Bayesian rRNA Classifier Version 2.11 (training set 16). The two reference datasets provided equivalent taxonomic assignments and confirmed the identity assigned to the 16S V4 rRNA sequences. One of the most dominant ESVs belonging to the Enterobacteriaceae family was unidentified at the genus level but isolation of that strain followed by Sanger sequencing on the full-length 16S rRNA gene revealed that it belongs to the genus *Klebsiella*. We therefore assigned that ESV to *Klebsiella*. The two *Klebsiella* isolates display different morphologies on glucose agarose plates and an indole test (Remel Kovacs Indole Reagent, #R21227) revealed that one of the isolates is indole positive while the other isolate is indole negative. Based on this distinction, we decided to refer to the two Klebsiella as Klebsiella positive (*Kp*) and Klebsiella negative (*Km*).

### Mapping isolates to amplicon sequencing data

To match our isolates from Sanger sequencing (full-length 16S rDNA sequence) to the amplicon sequencing data (ESVs) of the communities, we performed a pairwise alignment using the function *pairwiseAlignment* from the R package *Biostrings* (Pagès et al. 2017), with alignment type set to “*local*”. For each isolate in a community, we aligned its full-length Sanger sequence with all possible ESVs from the same community and obtained the reported alignment scores. Sanger-ESV alignment with highest alignment score was picked. If two Sanger sequences were matched to one ESV, the one with lower alignment score was dropped (19 of 73 isolate Sanger sequences were dropped). In the 54 pairwise alignments, the shortest consensus length is 234 base pairs, with 45 full matches, eight one base pair mismatches, and one two base pair mismatches.

### Bottom-up invasion experiment

We performed an invasion experiment between Klebsiella (*Kp*) (resident) and *Pseudomonas* and/or *Alcaligenes* (invaders) either alone (mono-invasion) or together (co-invasion). Prior to the start of the invasion experiment, the three strains were grown from frozen glycerol stocks alone into LB-agarose plates. For each strain, colonies were re-suspended into 1× M9 (without glucose) and normalized to an OD620 of 0.1. The normalized *A* and *P* stocks were then serially diluted independently 10-fold four times from 10^−1^ to 10^−4^. Note that here we refer to OD620 of 0.1 as the baseline OD (10^0^). For the co-invasion assays, *Alcaligenes* and *Pseudomonas* were mixed together for all five *A* and *P* dilutions (10^0^ to 10^−4^) generating 25 different *A-P* initial density combinations. Competitions were started by mixing 2uL of *Kp* with 2uL of the *A*:*P* mixtures (co-invasion) or 2uL of A or P monocultures (monoinvasion) at the 5 different dilutions into 500uL of M9 + glucose. In total, 36 invasion scenarios with different initial frequencies and densities of *A* and/or *P* (25 co-invasions, 10 mono-invasions, and *Kp* alone) were investigated in duplicate, setting the initial position of the population in the phase portrait shown in **Fig 3**. Strains were grown for 48h without shaking at 30C and then diluted 1:125 into fresh medium, and this growth-dilution cycle was repeated for 12 transfers. The relative abundance of each strain was estimated by plating (Colony-Forming Units) on LB-agarose plates.

### Phase diagram and separatrix

The phase diagram was obtained by analysis of the outcome of the bottom-up invasion experiment described above and shown in **Fig. 3** and **Fig. S12**. The ‘transition’ region was determined as follow. First, we identified the ‘flickering’ region of the *A-P* initial frequency space where the outcome either changed (e.g. from coexistence (gray) to A dominated state (yellow)) or remained gray at any point during one of the 3 transfers (T3, T8, T12) and in one or both of the 2 replicates analysed. The basin boundary of *Alcaligenes* was determined by taking, for each initial frequency, the mid-points between the initial frequencies inside the basin of attraction of *Alcaligenes* and inside the ‘flickering’ region that are closest to the transition line. Similarly, the basin boundary of *Pseudomonas* was determined by taking, for each initial frequency, the mid-points between the initial frequencies inside the basin of attraction of *Pseudomonas* and inside the ‘flickering’ region that are closest to the transition line. The separatrix shows the midline between the two boundaries. In Fig. 3A, the datapoints where the relative abundance is below the detection threshold are arbitrarily set to a value of −4.33.

## Acknowledgements

We want to thank Alicia Sanchez-Gorostiaga and Josh Goldford for helpful discussions, and Jackie Folmar for assistance with platting. We thank Sven Even Borgos and Juan Nogales for providing us with SBML versions of *Pseudomonas* genome-scale metabolic models. The funding for this work partly results from a Scialog Program sponsored jointly by Research Corporation for Science Advancement and the Gordon and Betty Moore Foundation through grants to Yale University (AS). This work was also supported by a young investigator award from the Human Frontier Science Program (RGY0077/2016) and by the National Institutes of Health through grant 1R35 GM133467-01 to AS. MRG was additionally supported by a Donnelley Postdoctoral Fellowship from the Yale Institute for Biospheric Sciences.

