## Supplementary Material for "Metabolic rules of microbial community assembly"

This file includes:

- Supplementary figures **(Fig. S1 –Fig. S17)**
- Supplementary tables **(Table S1- Table S2)**

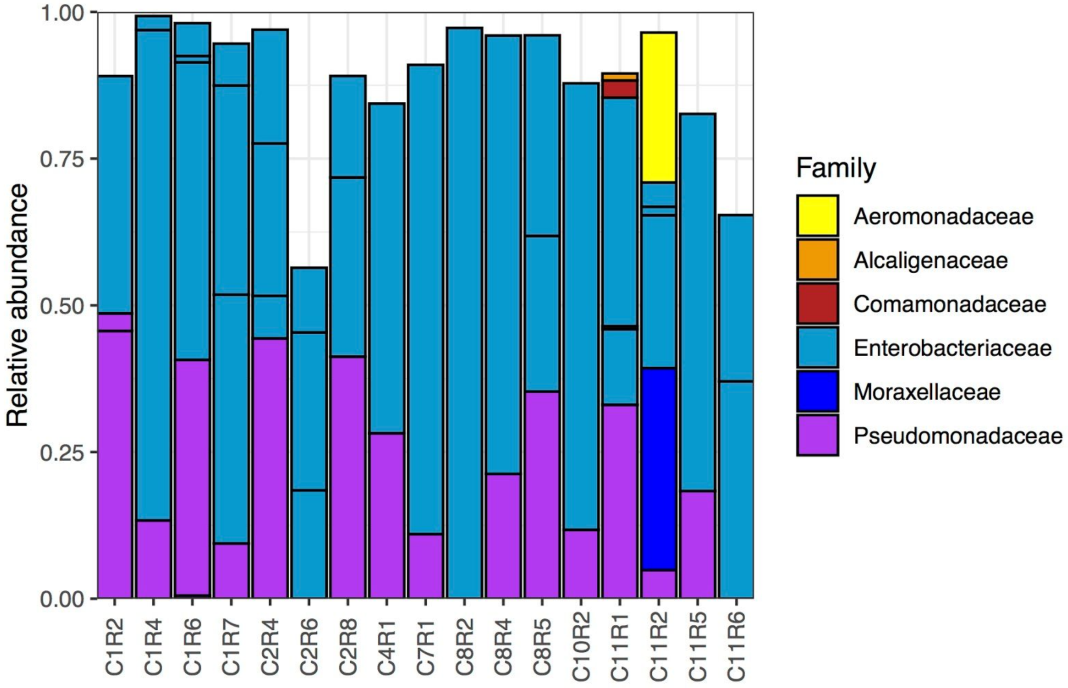


**Fig. S1. The strains isolated represented most of the taxonomic composition of the 17 communities from where they were collected.** N=73 strains isolated from 17 communities (CxRy) covering 88.8% of the taxonomic composition (Methods).

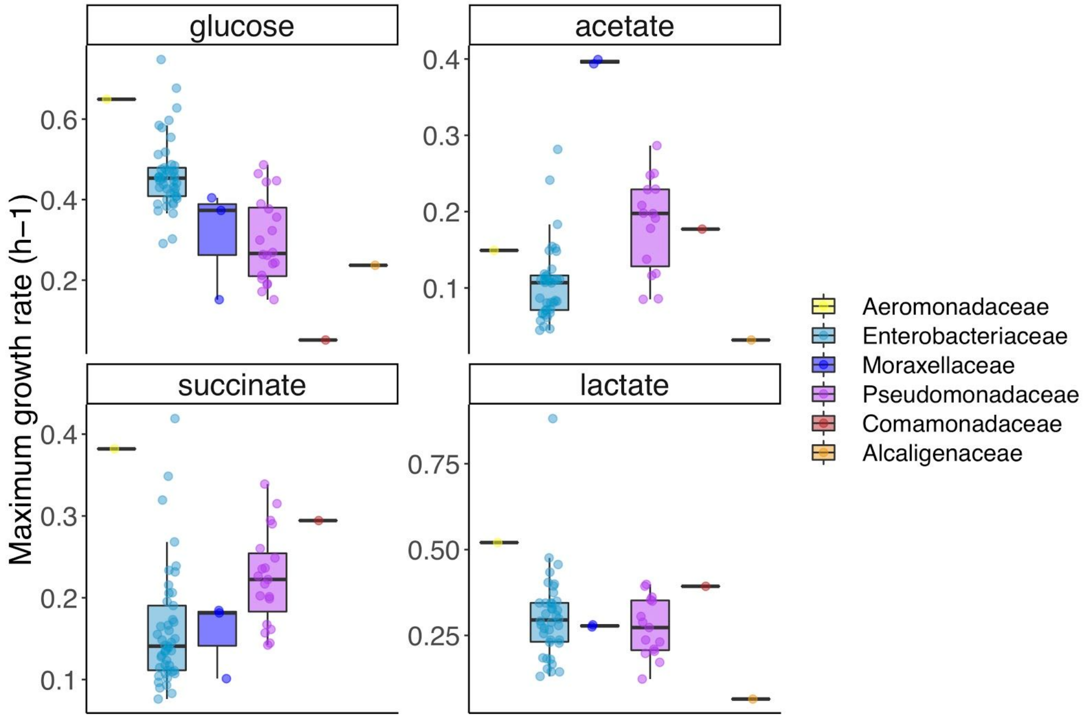


**Fig. S2. Family-level variation in maximum growth rate.** Maximum growth rate of strains belonging to different families when grown in minimal media supplemented with either glucose, acetate, succinate, or lactate as single carbon source. N=73 strains.

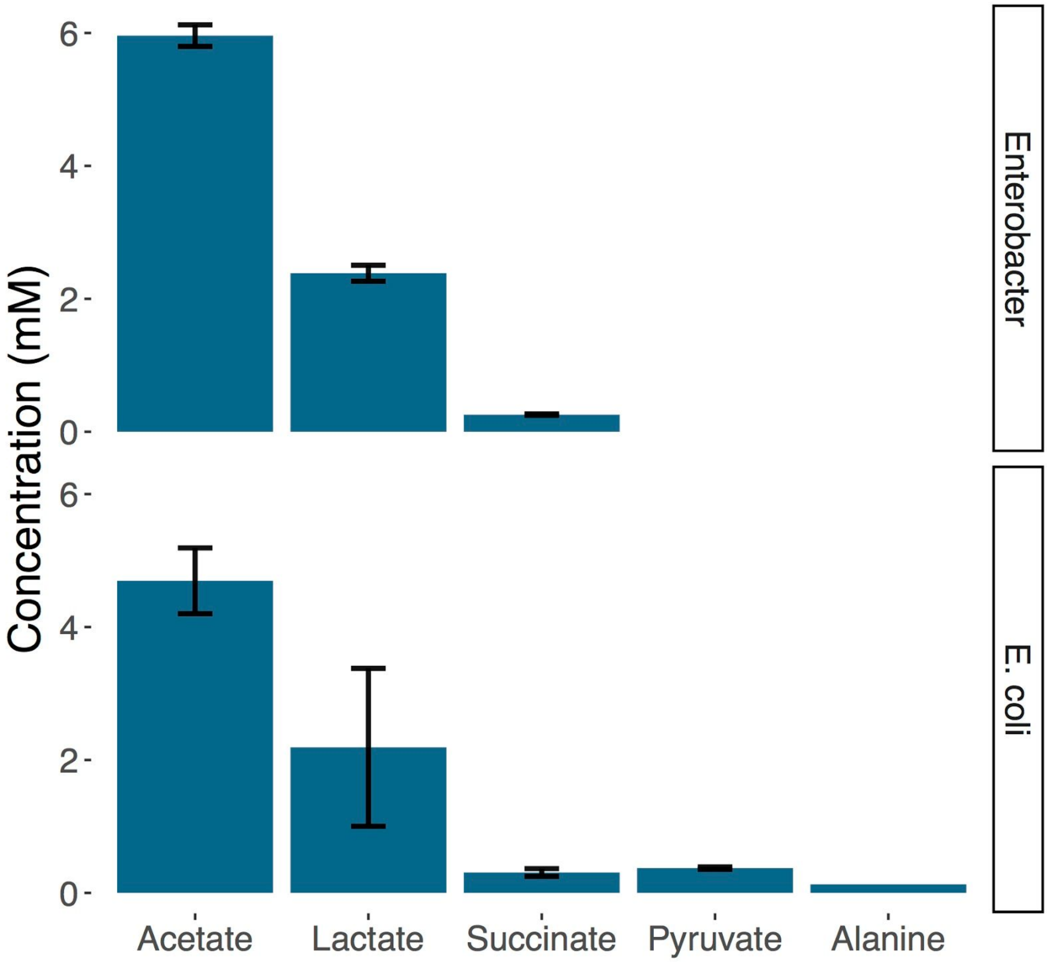


**Fig. S3. Concentration of metabolites secreted by *E. coli* and *Enterobacter* growing on glucose.** Metabolites found in the supernatant of each monoculture after 28h of growth in minimal glucose media. Metabolites were quantified through LC-MS. Shown are the metabolites with concentrations above 0.1 mM. Shown is the mean ± sd of two replicates.

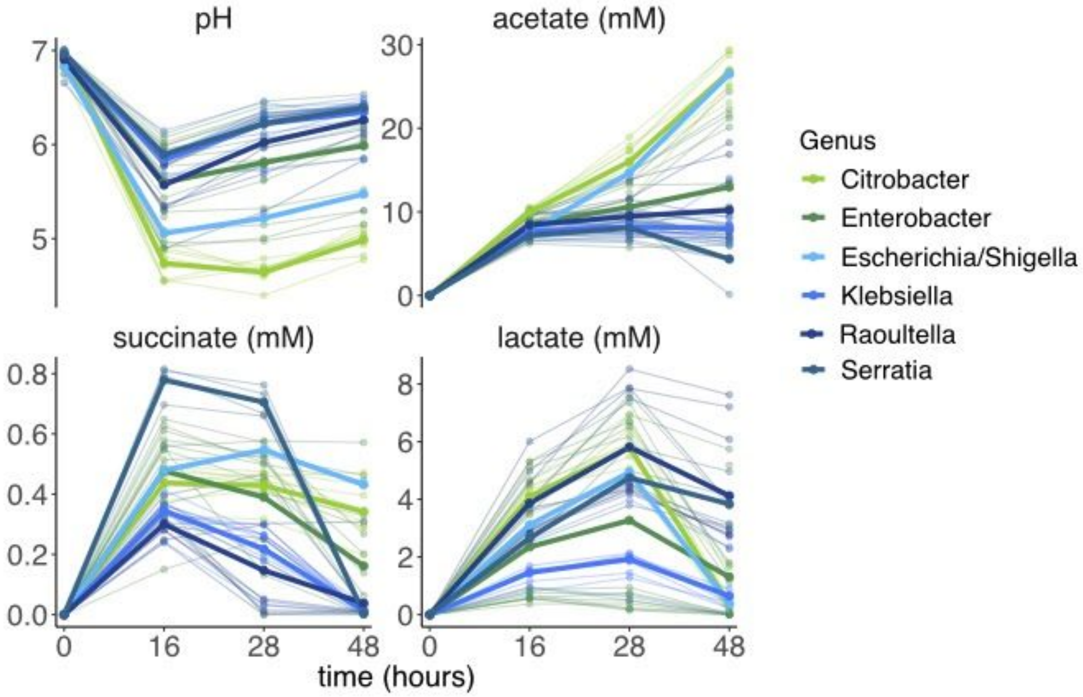


**Fig. S4. Genus-level variation in the amount of organic acids excreted by Enterobacteriaceae.** pH and concentrations of acetate, succinate, and lactate in the supernatant of strains grown in minimal media with glucose. Supernatant collected at different timepoints during a 48h growth cycle. The thick line shows the mean of each genus (N=73 strains).

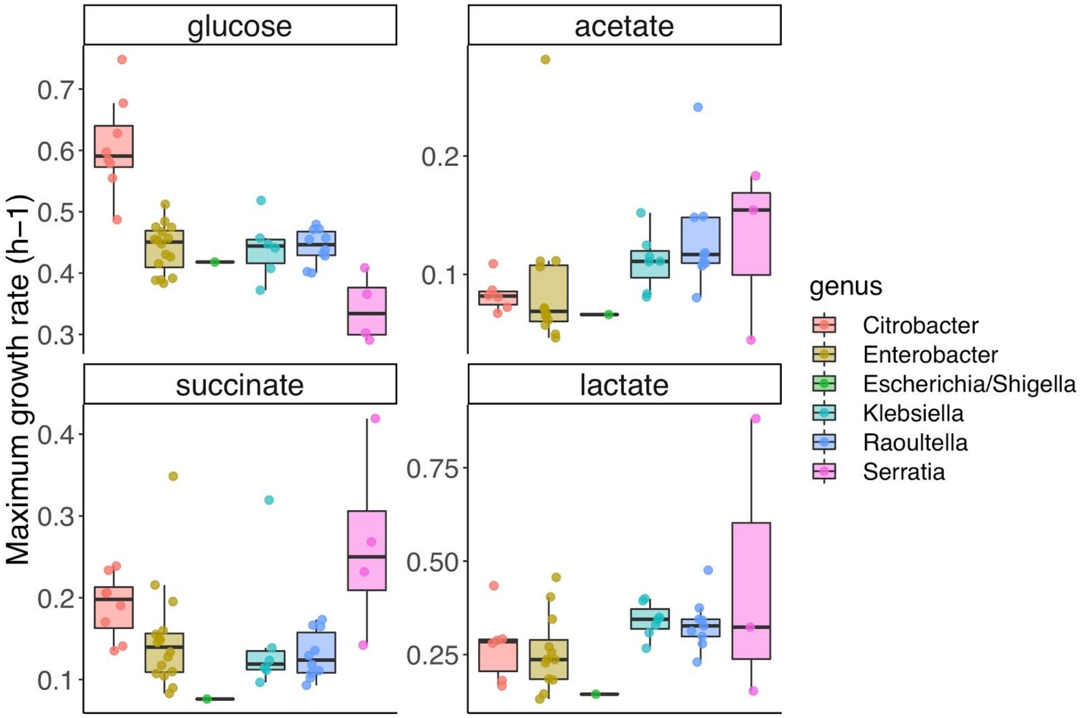


**Fig. S5. Genus-level variation in maximum growth rate.** Maximum growth rate of different strains belonging to the Enterobacteriaceae family when grown in minimal media supplemented with either glucose, acetate, succinate, or lactate as single carbon source.

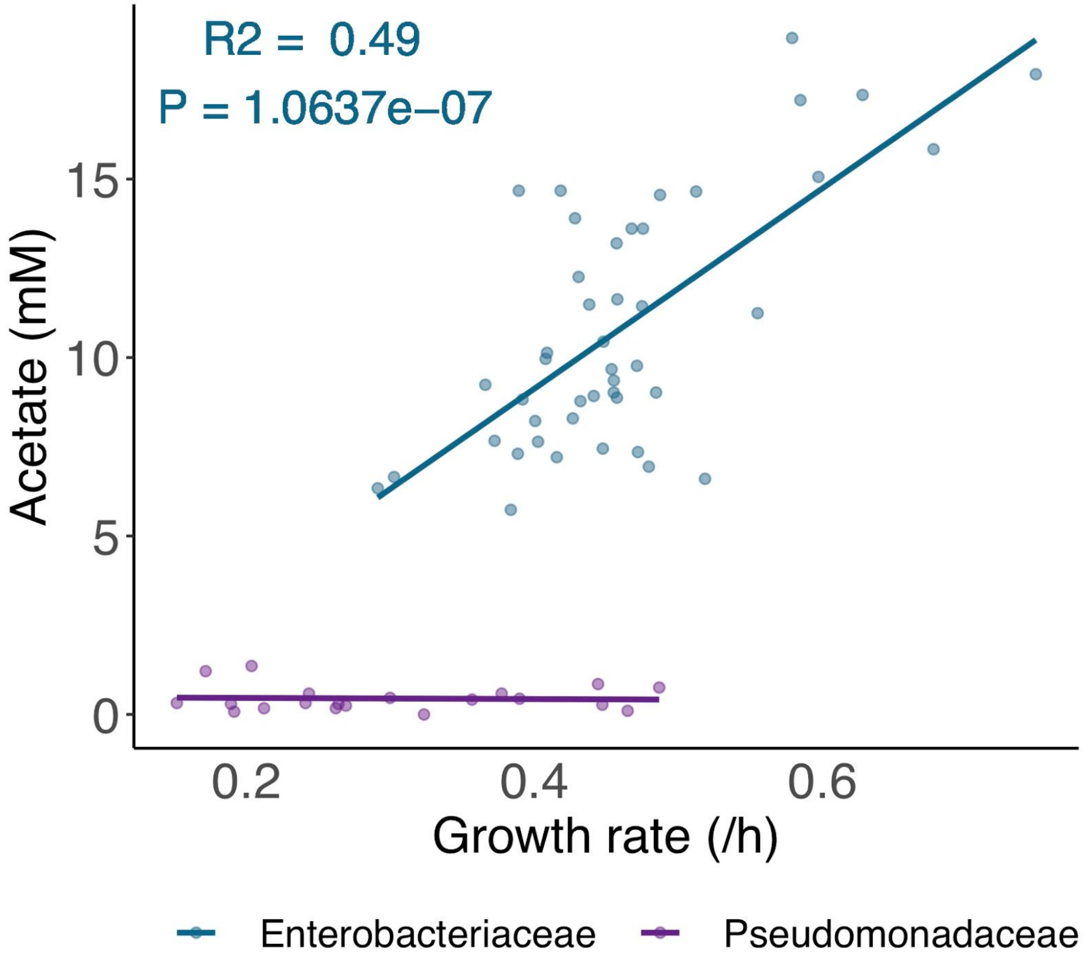


**Fig. S6. The amount of acetate excreted by the Enterobacteriaceae species correlates with their maximum growth rate in glucose.** Maximum growth rate of different strains belonging to the Enterobacteriaceae (N=47) and Pseudomonadaceae family (N=20) when grown in minimal glucose media. Acetate concentration in the supernatant after 28h of monoculture growth in minimal glucose media. The lines are the linear regression lines for each family.

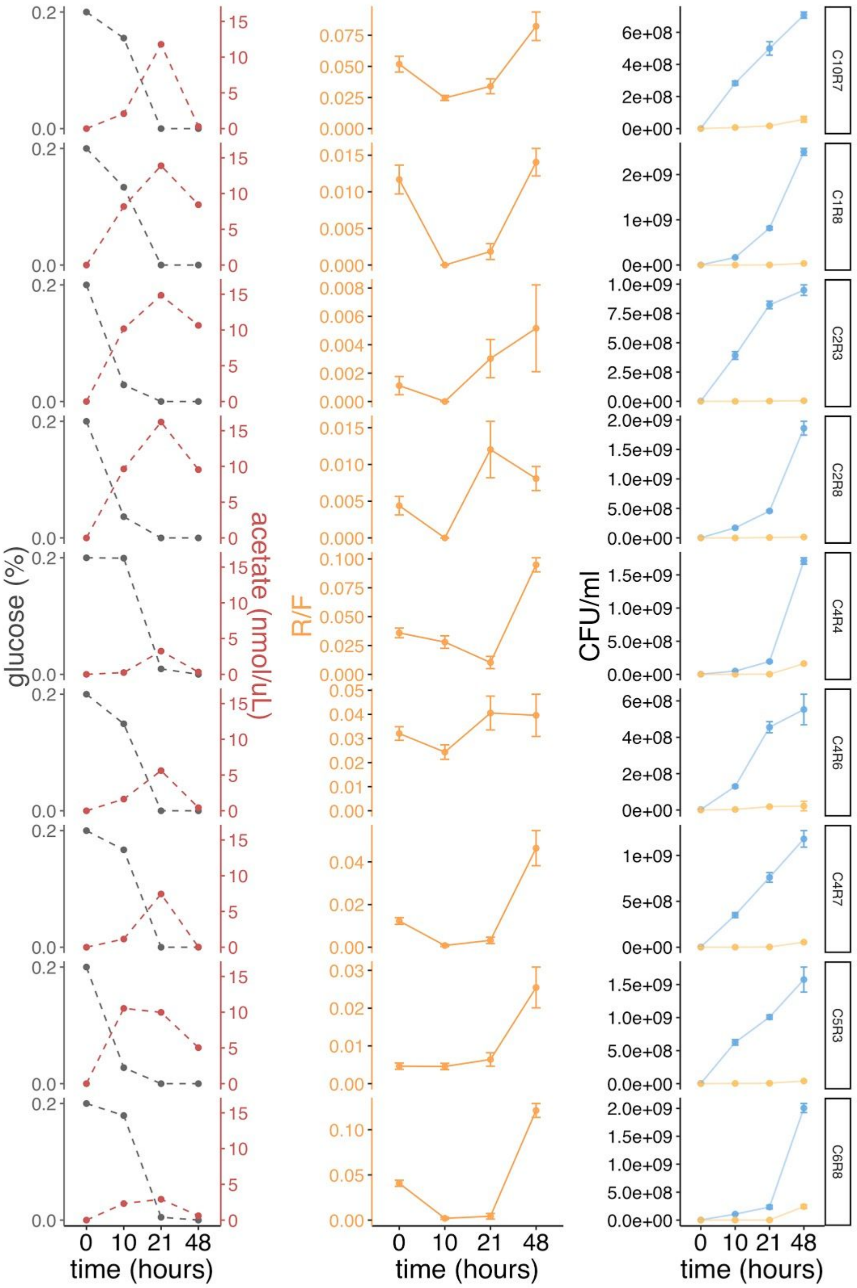

glucose (%)
acetate (nmol/uL)
R/F
CFU/ml
time (hours)
C10R7
C1R8
C2R3
C2R8
C4R4
C4R6
C4R7
C5R3
C6R8

**Fig. S7. Growth of fermenters is associated with glucose depletion, whereas growth of respirers is associated with acetate depletion.** We thawed whole communities and revived them in glucose minimal media for three passages. We selected a subset of communities where fermenters and respirers coexist after three serial transfers (N=9 communities from 6 different inocula where CxRy corresponds to inoculum x, replicate y) and measured their R/F ratio, and concentrations of glucose and acetate at different timepoints during a 48h growth cycle (0h, 10h, 21h and 48h). The R/F ratio represents the mean $\pm$ sd of the CFU ratios calculated using a bootstrap method (N=1000 replicates). For three of the communities (C1R8, C2R3, and C2R8), there were no detectable R colonies on the plates, so we set R/F to 0. To make sure that the observed decrease in R/F ratio from 0h to 10h is not due to differences in the total number of CFUs across timepoints, we confirmed that the total CFU count at 10h is larger than the total CFU count at 0h. C10R7 corresponds to the community shown in **Fig 1C**.

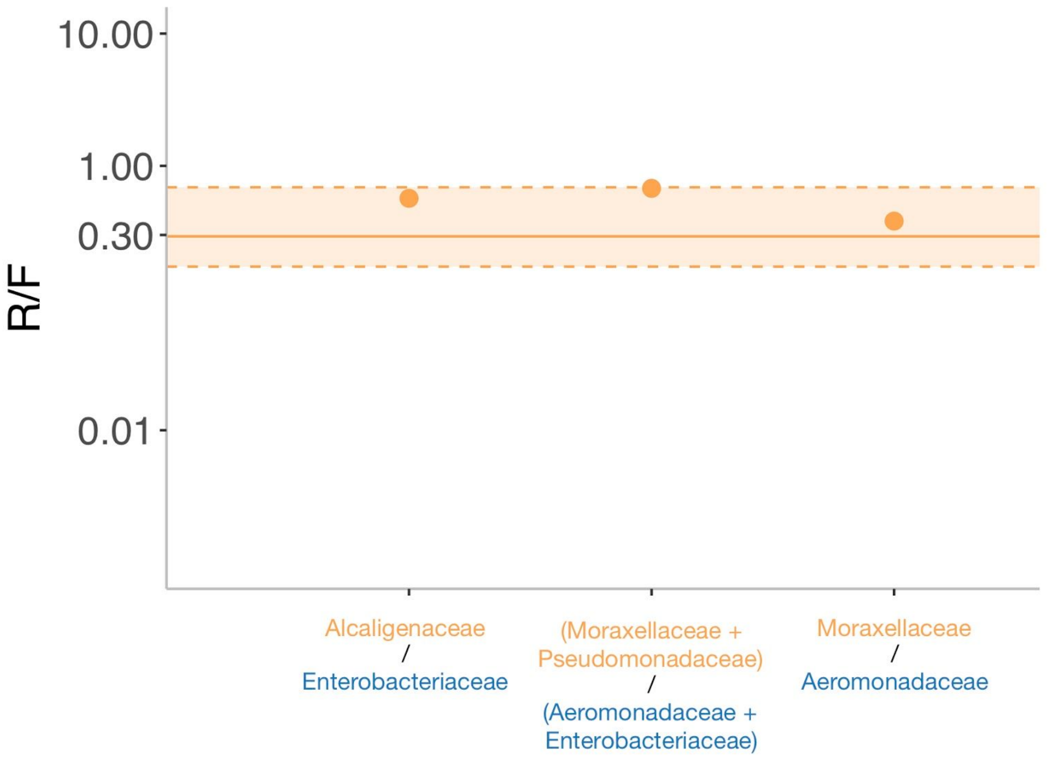


**Fig. S8. Communities dominated by other respirator and fermenter families exhibit similar R/F ratios.** Shown is the median R/F ratio. The Alcaligenaceae/ Enterobacteriaceae communities were taken from **Fig. 2B** (state dominated by Enterobacteriaceae and Alcaligenaceae as dominant F and R respectively; N=65, median =0.56, Q1=0.41, Q3=0.66); the (Moraxellaceae+Pseudomonadaceae/Aeromonadaceae+Enterobacteriaceae) community corresponds to one community (C11R2) in Goldford et al. (2018) that is not dominated by *Pseudomonas* and Enterobacteriaceae (N=1, median=0.68), and the Moraxellaceae/ Aeromonadaceae communities were from an experiment where one community (12 replicates) was passaged for 10 transfers in 5mL of M9+glu inside a 50mL Falcon tube under the same other experimental conditions as before (N=12, median=0.36, Q1=0.28, Q3=0.51). The horizontal line shows the median R/F ratio of the glucose communities presented in **Fig. 1A** (R/F=0.29) and the dashed lines show the interquartile range (Q1=0.17, Q3=0.69).

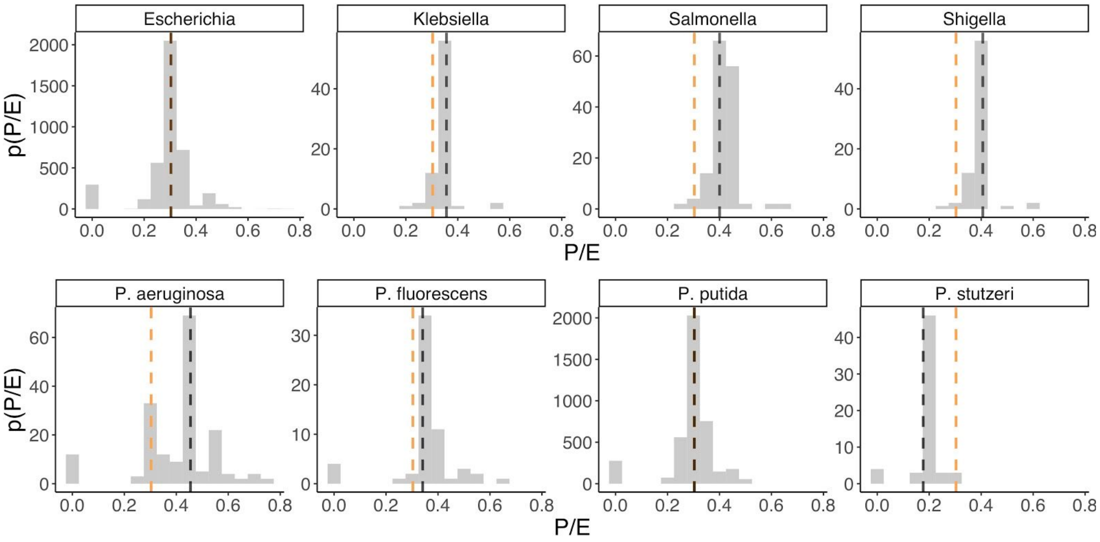


**Fig. S9. Probability distribution of the FBA predicted P/E ratio for different pairs of Enterobacteriaceae -Pseudomonas models.** Distributions are shown separately for each genus of Enterobacteriaceae, or species of *Pseudomonas.* The yellow line shows the median of all Pseudomonas/ Enterobacteriaceae pairs (P/E=0.3) and the black line shows the median for the pairs shown in the respective facet.

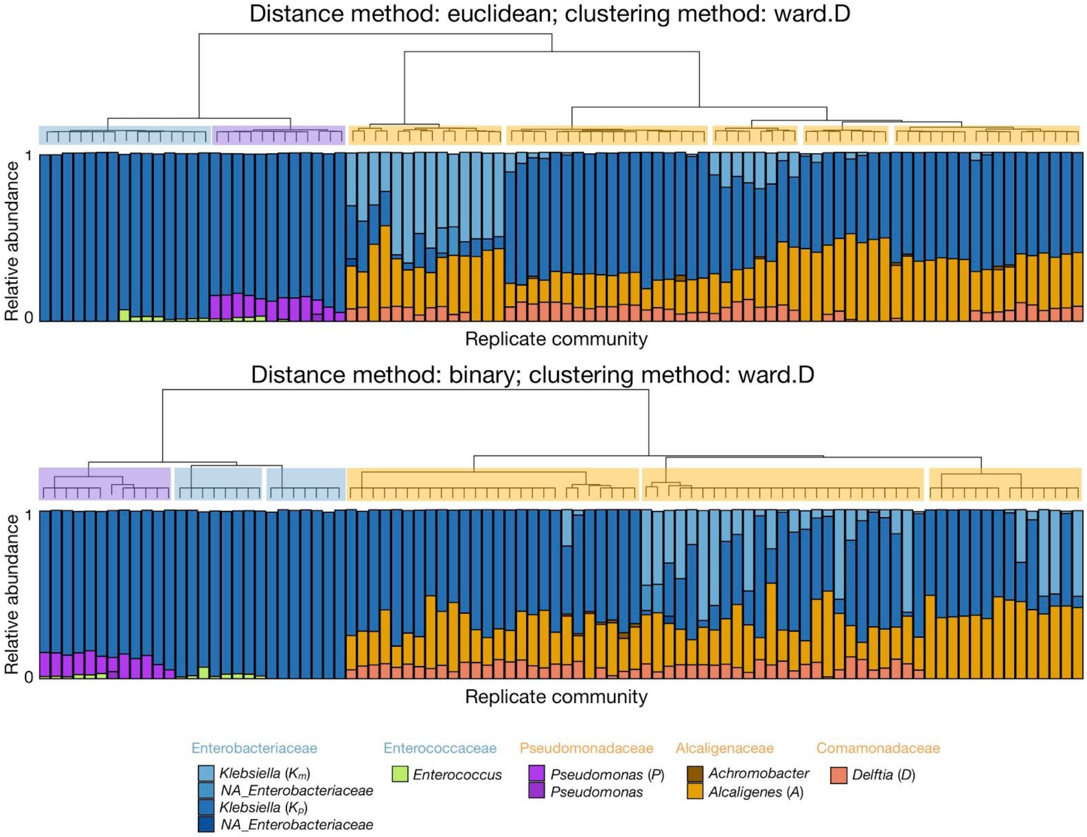


**Fig. S10. Detection of alternative states in community composition.** Taxonomic profile of communities shown at the exact sequence variant (ESV) level (one color per ESV) with corresponding genus and family level assignments. Only the ESVs with a minimum relative abundance > 0.01 are shown. The communities are sorted according to the order obtained from the hierarchical tree (dendrogram) of the relative abundance of all ESVs using the "ward.D" clustering method and either the "euclidean" (top) or binary (bottom) distance (*hclust* R function). Six alternative states in community composition (clusters) obtained using the *cutree* function in R to cut the tree generated by *hclust* are coloured as in **Fig. 2B**.

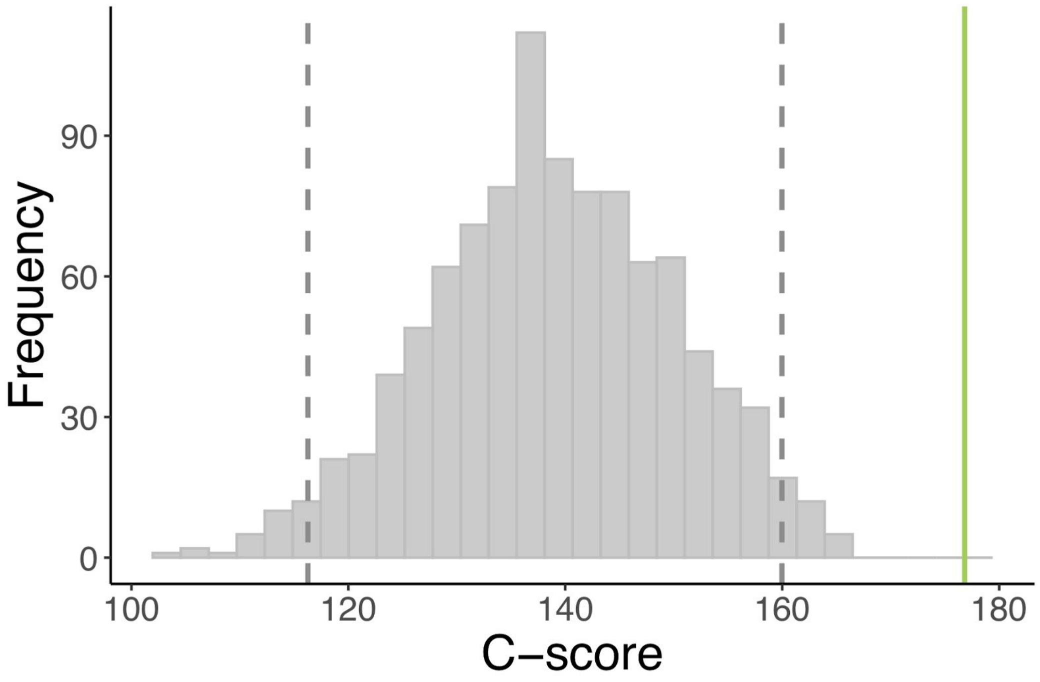


**Fig. S11. Interactions between species are non-random, and are instead highly segregated.** Distribution of C-scores expected using a null model for the assembled communities. The observed C-score for our communities (green line) is significantly higher than we would expect by random chance given the species in the communities ($p_{obs>null}$ <0.001), suggesting that there is a high degree of segregation between species. The gray dashed lines indicate the lower and upper 95% confidence interval (two-tailed), respectively. Only taxa with a relative abundance > 0.01 were considered.

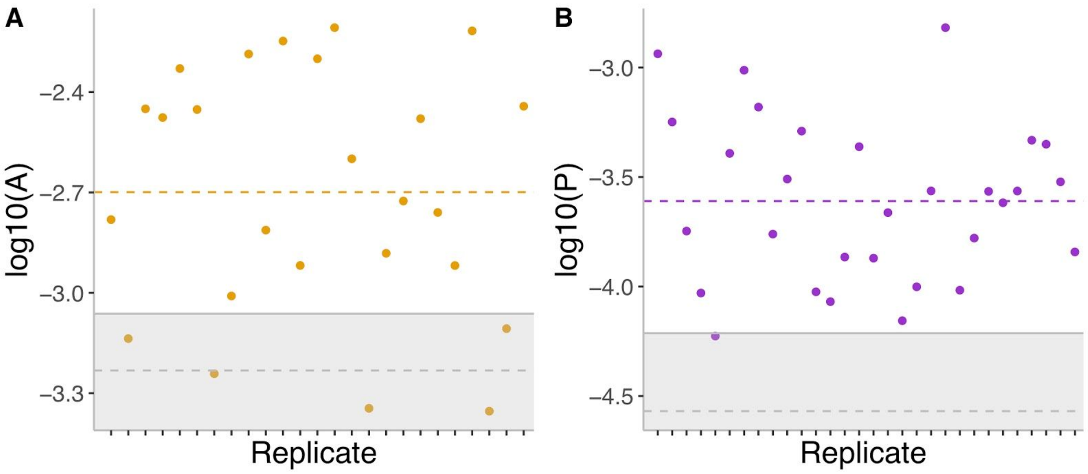


**Fig. S12. Relative abundance of *Alcaligenes* and *Pseudomonas* for communities with low *Alcaligenes* (A) and with low *Pseudomonas* (B).** The dashed coloured line shows the mean. The dashed gray line is the mean of *A* and *P*, respectively, in the controls (N=22, Methods) and the filled line is the 95% confidence interval. To rule out the hypothesis of random sampling, only the communities that pass the 95% confidence interval test (that is, above the shaded area) were included in the calculation of communities that contain both *A* and *P* after 18 transfers.

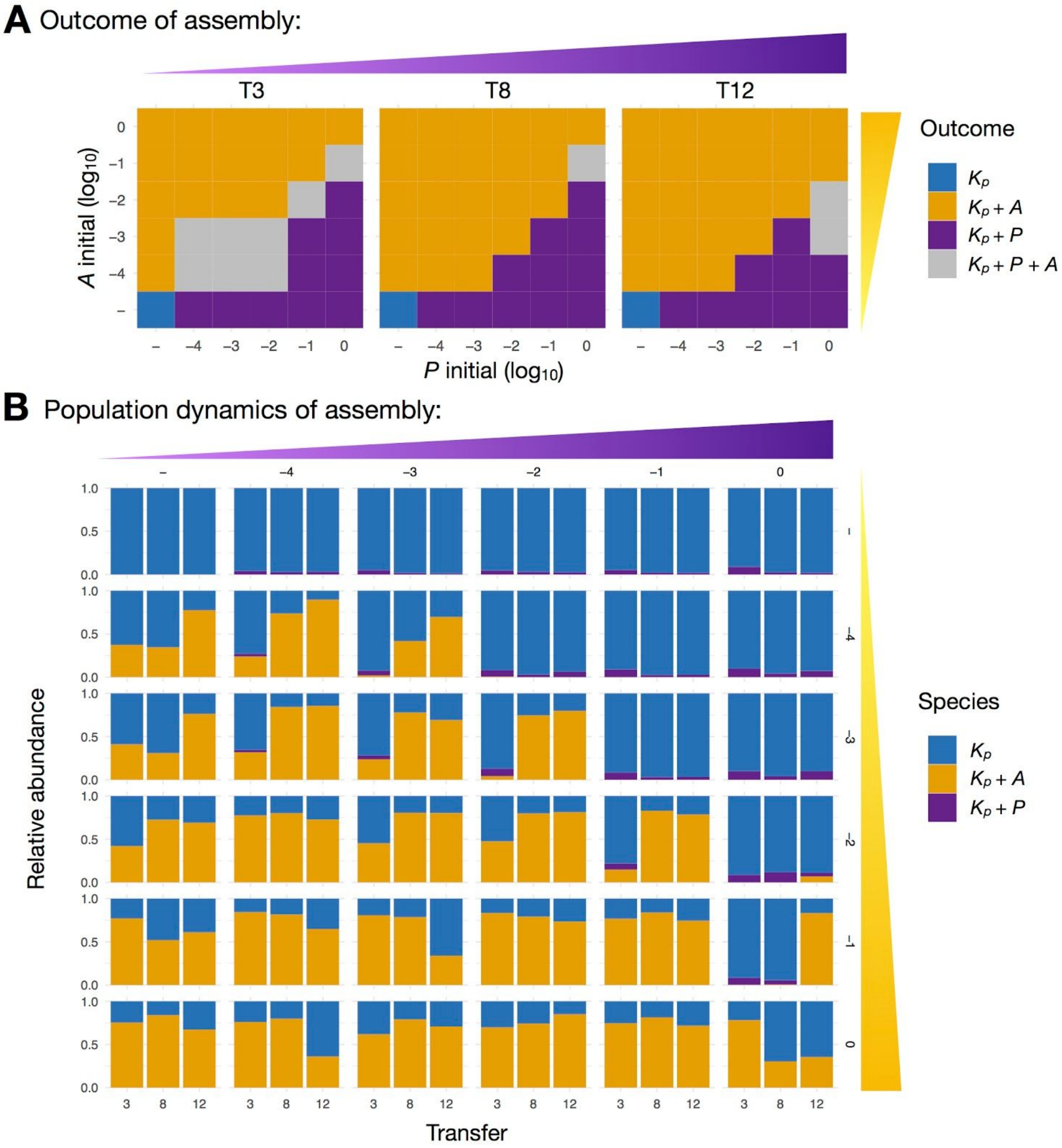


**Fig. S13. Outcome and population dynamics of assembly of reconstituted communities for second replicate.** Results for replicate 1 are shown in **Fig. 3** and **Fig. S14**.

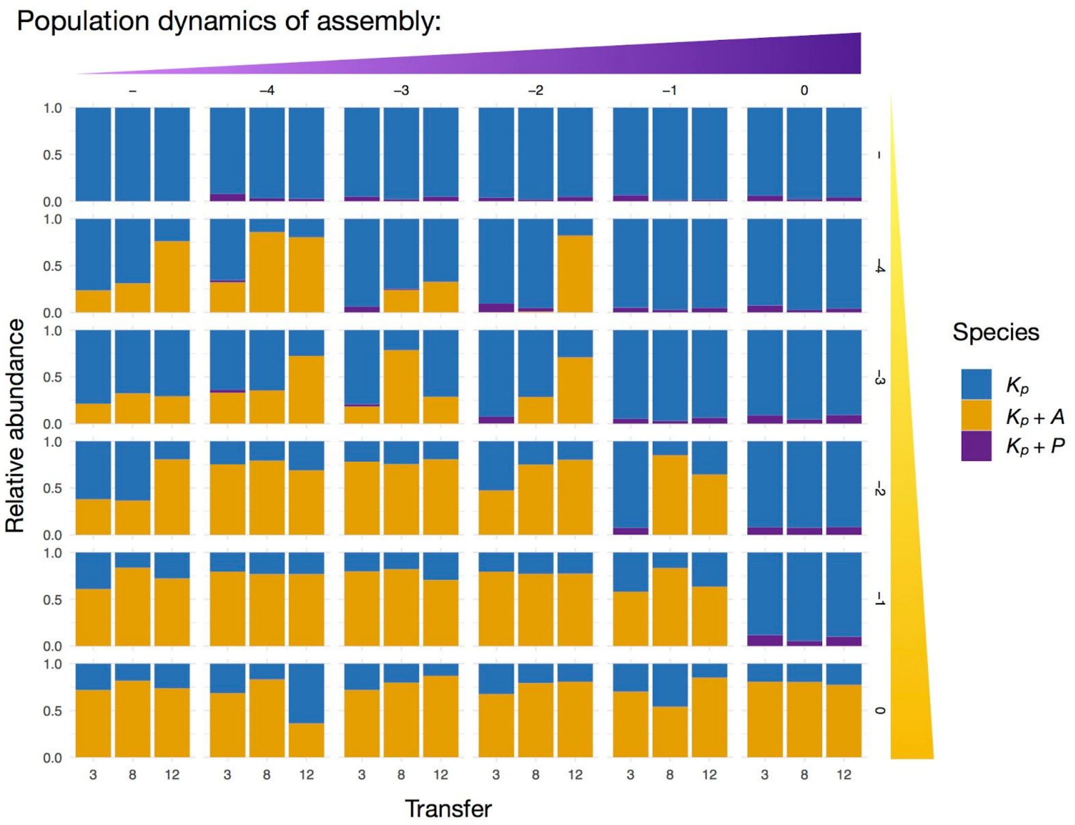


**Fig. S14. *Alcaligenes* and *Klebsiella* or *Pseudomonas* and *Klebsiella* coexist in reconstituted communities after 12 transfers.** Shown are the population dynamics of each species when started under different initial densities/frequencies of *Alcaligenes* (*A*) and/or *Pseudomonas* (*P*) for the replicate shown in the main text (**Fig. 3**). *P* increases from left to right whereas *A* increases from bottom to top. Outcome of assembly is shown in **Fig. 3**.

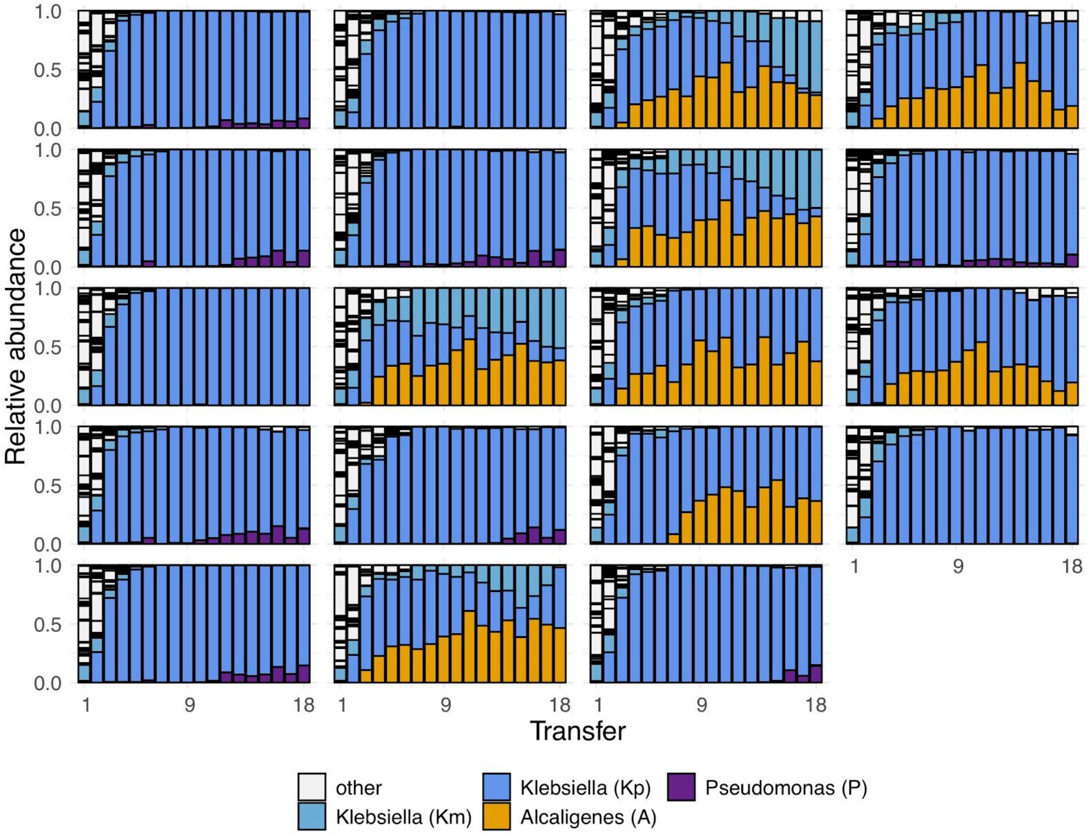


**Fig. S15. Temporal dynamics of replicate communities assembled in glucose media.** Replicate communities were all started from the same inoculum, and serially diluted in minimal media with glucose every 48h for a total of 18 growth-dilution cycles. Only the top four dominant ESVs at transfer 18 are coloured, other ESVs are shown in gray.

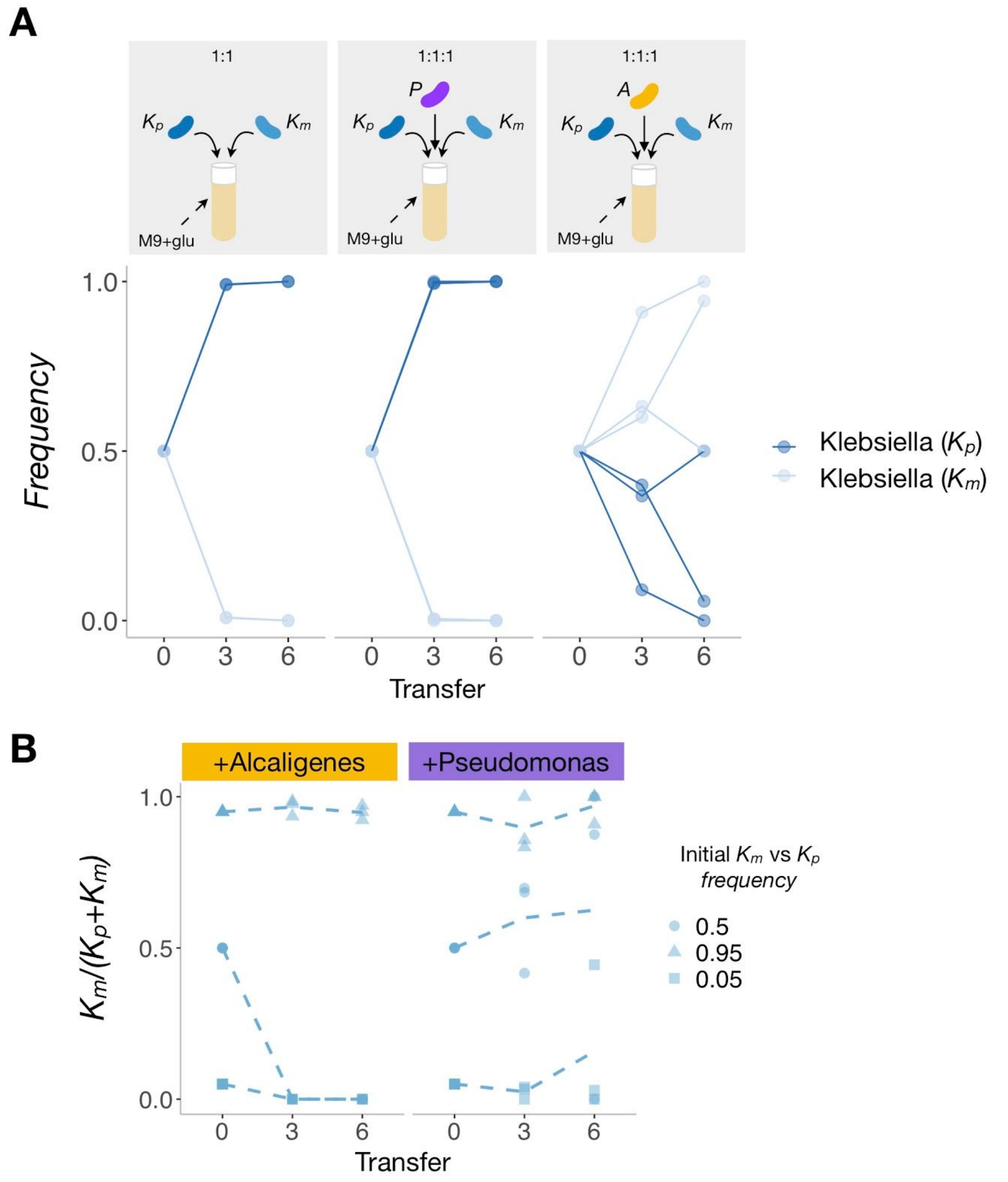


**Fig. S16. *Alcaligenes* buffers coexistence between the two dominant *Klebsiella* strains. A.** *Kp* and *Km* were inoculated in duo (1:1) and trio (1:1:1) communities with either *Alcaligenes* or *Pseudomonas*, and, **B** in trio with different initial frequencies (0.05, 0.5, or 0.95), and then passaged for six growth-dilution cycles (12 days) in M9+glucose. Communities were plated and CFUs were counted in LB agarose (to distinguish *A* and *P* from *K*) and in M9+glucose agarose (to distinguish *Kp* from *Km*). There are 2 biological replicates for the duos and 3 replicates for the trios.

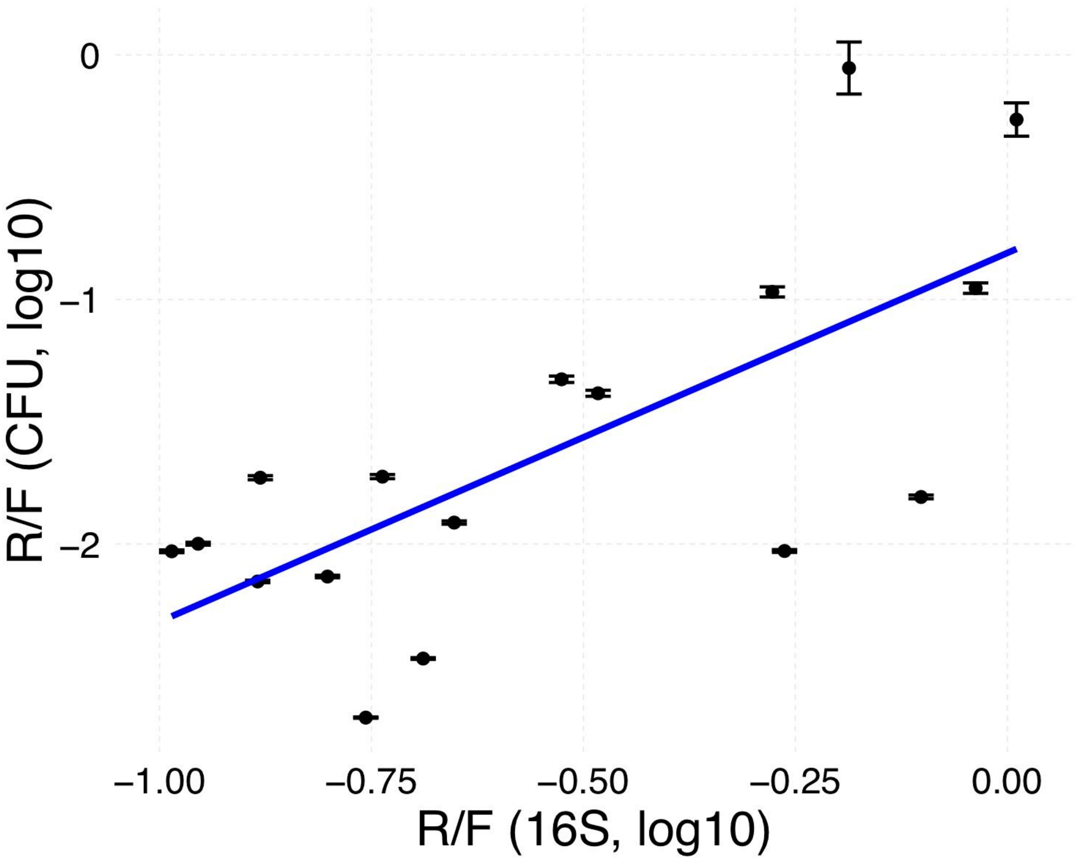


**Fig. S17. Positive correlation between the R/F ratio measured by CFU and by 16S sequencing.** Log-log Respirator:Fermenter ratio in glucose communities (T12) from Goldfored et al. (2018). 16S data from Goldford et al (2018). To calculate CFUs ratio, 24 communities were thawed from frozen stock and CFUs were counted on chromogenic agar. White colonies were counted as R and blue/purple colonies were counted as F. Shown are communities with at least one colony of R and F (n=17). For each community, dots show the mean ± SD of CFU ratios calculated using a bootstrap method (N=1000 bootstrap estimates from resampling with replacement). Blue line is the regression line with slope= 1.51 and intercept -0.81 ($R^2$=0.49).

**Table S1. Sensitivity analysis on the parameters used for the CAFBA simulations for the *E. coli* (iJO1366) and *P. putida* (iJN1463) metabolic models**

| Parameter | Fold change | Value | P/E |
|---|---|---|---|
| $\phi_{\max}$ | Double | 0.968 | 0.350 |
| $\phi_{\max}$ | Default | 0.484 | 0.357 |
| $\phi_{\max}$ | Half | 0.242 | 0.370 |
| $w_E$ | Double | 0.001770326 | 0.365 |
| $w_E$ | Default | 0.000885163 | 0.357 |
| $w_E$ | Half | 0.000442581 | 0.353 |
| $w_R$ | Double | 0.338 | 0.361 |
| $w_R$ | Default | 0.169 | 0.357 |
| $w_R$ | Half | 0.0845 | 0.355 |
| *P. putida* limitation | TRUE | N/A | 0.355 |
| *P. putida* limitation | FALSE | N/A | 0.357 |

**Table S2. Glucose fermentation profile of dominant families**

| Family | Fermentation profile |
|---|---|
| Enterobacteriaceae | Fermenter |
| Enterococcaceae | Fermenter |
| Aeromonadaceae | Fermenter |
| Pseudomonadaceae | Respirator |
| Alcaligenaceae | Respirator |
| Moraxellaceae | Respirator |
| Xanthomonadaceae | Respirator |

### Supplementary Methods for

### Metabolic rules of microbial community assembly

Sylvie Estrela, Jean C. C. Vila, Nanxi Lu, Djordje Bajic, Maria Rebolleda-Gomez, Chang-Yu Chang, and Alvaro Sanchez

This file includes Supplementary Methods for:

- Constrained-Based Modelling
- Metabolic Model Datasets

#### Constrained-Based Modelling

CAFBA and FBA simulations were performed using the COBRApy package. CAFBA is an extension of FBA which explicitly incorporates a global constraint on proteome allocation (Mori et al. 2016). Unlike FBA, CAFBA correctly predicts the secretion of acetate at high growth rates (Basan et al. 2015) . The exact notation of the constraint is as follows:

$$\max_{\mathrm{v}\in\mathcal{F}} \mu \quad \mathrm{subj.\,to} \quad w_C J_C + w_R \mu + \sum_{i\in E} w_i |v_i| = \phi_{\max}.$$

To simulate *E. coli* growth on excess glucose we follow the approach taken by Mori et al. (2016). We set $\phi_{\max}$ and $w_R$ to their *E. coli*-specific empirical values of 0.484 and 0.169/h. The cost $w_i$ of all reactions in the E-sector is set to some constant $w_E$ such that a maximum achievable growth rate on glucose ($w_C \rightarrow 0$) is 1/h. Because glucose is in excess $w_C$ =0.0 and glucose uptake is unbounded. We also set the following nutrients to unbounded: ca2_e, cbl1_e, cl_e, co2_e, cobalt2_e, cu2_e, fe2_e, fe3_e, h_e, h2o_e, k_e, mg2_e, mn2_e, mobd_e, na1_e, nh4_e, ni2_e, pi_e, sel_e, slnt_e, so4_e, tungs_e, zn2_e. As in Mori et al we silence the glucose dehydrogenase reactions (GLCDe and GLCDpp). Growth is optimized using cobrapy's optimize function and we record the flux through the objective function (biomass/hr) as well as the fluxes through the exchange reactions.

To simulate *P. putida* growth on secreted metabolites we use Flux Balance Analysis (FBA). We first set all metabolites to be unavailable (lower bound of 0) excluding the same set of inorganic ions as before. For each metabolite secreted by *E. coli* we set the lower bound on P. putida's uptake to match the predicted secretion rate. Growth is then optimized using cobrapy's *optimize* function and we record the flux of the biomass reaction (biomass/h). The predicted P/E ratio is the flux through biomass reaction of the *P. putida* metabolic model divided by the flux through the biomass reaction of the *E. coli* metabolic model. For iJO1366 and iJN1463 we get a PE ratio of 0.356. We find that this value is extremely robust to the exact CAFBA parameters used. The parameters $w_E$, $\phi_{\max}$, and $w_r$ can all be halved or doubled without substantially changing the predicted P/E ratio **(Table S1)**. Furthermore, whilst we simulated *P. putida* using conventional FBA, we note that if we do impose the CAFBA constraints on the *P. putida* metabolic model using the same approach as for *E. coli* (modeling *P. putida* as growing on *E. coli* secretions as though they were in excess), we obtain a P/E ratio of 0.355 which is extremely close to the value reported in the main text. We repeat the above analysis for every pair of Enterobacteriaceae and Pseudomonadaceae metabolic model in a manually compiled model collection (see below). This was used to obtain a distribution of predicted P/E ratios.

We make note of minor technical differences between our simulations and those of (Mori et al. 2016) in how we determine which reactions belong to the E-sector. Here the E-sector is defined as including all enzyme catalyzed reactions except for transporters and exchanges. Unlike in Mori et al where E-sector identity was determined using SBML subsystem assignment (which are not available for all metabolic models) we define transporters using reaction stoichiometry and metabolite compartments which are available for all metabolic models and can be inferred from metabolite IDs. Ignoring sodium and hydrogen ions, a transporter is defined as any reaction that consumes at most one metabolite in one compartment and produces at most one metabolite in another compartment (cytoplasm, periplasm and extracellular). This includes all common transporter types (symporters, diffusion reaction etcs) whilst excluding membrane associated reactions such as those involved in oxidative phosphorylation (which are part of the E-sector). For the iJR904 model used by Mori et al. (2016), this approach gives us $w_E$ for all reactions in the E-sector of 0.00084 which is extremely similar to the 0.00083 they obtained. For the iJO1366 model this approach gives us a $w_E$ of 0.000885.

**Metabolic Model Datasets**

To compile a collection of Enterobacteriaceae metabolic models we downloaded every available metabolic model from the biggs database (version 1.6). We used every Enterobacteriaceae models, except for those that could not grow on minimal glucose. This gave us 59 metabolic models (55 *Escherichia*, 1 *Klebsiella*, 2 *Salmonella*, and 1 *Shigella*).

Because only 2 *Pseudomonas* metabolic models are available in the biggs database (and iJN746 is merely an older version of the iJN1463 metabolic model), we instead compile a library of 74 *Pseudomonas* metabolic models from the literature. Including iJN1643, 69 out of the 74 metabolic models were for different strains of *P. putida*. We constructed these models using the published gene orthology matrix and approach outlined in Table S5 of (Nogales et al. 2020). For each strain of *P. putida* we started with the iJN1463 metabolic model and removed genes and reactions that were missing. Genes with less than 80% percentage identity were considered to be missing. As in Nogales et al. (2020), we excluded strains with >300 genes missing. Gapfilling was then performed using Cobrapy's *gapfilling* module to identify the reaction from iJN1463 that would enable growth on minimal acetate. These reactions were then added back to create the strain specific model. To confirm that our results would apply to Pseudomonads other than *P. putida*, we also included three *P. aeruginosa* metabolic models (iMO1056, iPAE1146, iPAU1129) and as well as a *P. stutzeri* metabolic model (iPB890) and a *P. Fluorescens* metabolic model (iSB1139). **Fig 1D** shows the full distribution of predicted P/E values whilst in **Fig. S9** we break down the results by Enterobacteriaceae genus and *Pseudomonas* species.
